# Arbitrium phages can manipulate each other’s lysis - lysogeny decisions

**DOI:** 10.1101/2025.10.13.681967

**Authors:** R Manley, R Woodhams, J Bruce, E Smith, B Temperton, E. R. Westra

**Author notes:** Contributed equally (R Manley, R Woodhams).

## Abstract

Many viruses can switch between lytic replication and dormancy (or lysogeny). It was recently discovered that some viruses that infect bacteria (known as bacteriophage, or phage) employ peptide-based (“arbitrium”) communication systems to optimise their lysis/lysogeny switch: high peptide concentrations signal a lack of susceptible hosts and trigger lysogeny, while low peptide concentrations signal an abundance of uninfected hosts and prompt lysis. Here we demonstrate that Arbitrium-phages belonging to different species and genera can influence each others’ infection dynamics by secreting similar communication peptides, leading to early lysogenisation of the signal-receiving phage, and elevated fitness of the signal-emitting phage. Antagonistic coevolution between signal emitting and signal receiving phages to manipulate each other’s infection behaviours may explain the rapid diversification of arbitrium systems and their frequent horizontal exchange to escape the noise of cross talk.

## Introduction

Temperate phages can transmit both horizontally (lytic cycle) and vertically (lysogenic cycle). During the lytic cycle phages infect a host cell, replicate and kill the host, releasing virions into the environment ^1^; during the lysogenic cycle phages integrate into the host genome, persist as a prophage and replicate at the rate of the host ^2,3^. Importantly, a temperate phage can transition between these two states by reactivating its lytic cycle. When a temperate phage infects a bacterium as a free phage, or is integrated as a lysogen, it must ‘choose’ the infection strategy that optimises its fitness, which will depend on the availability of healthy, susceptible host cells to infect ^4^. If there are plenty of cells to infect it is optimal to be lytic, but when host cells are rare it is optimal to lysogenise the surviving hosts and remain dormant until host cell numbers increase. Optimising this decision is a key fitness determinant of temperate phage.

It was recently discovered that some phages that infect *Bacilli* bacteria use a peptide-based communication system, named the ‘arbitrium system’ to optimise their decision to lyse or lysogenise their host ^5^. The arbitrium system is analogous to bacterial quorum sensing and has been genetically and mechanistically characterised in the *Bacillus* phages Phi3T and SpBeta ^5–11^. The arbitrium system comprises a series of key genes (*aimP, aimR, aimX*) that enable the phage to produce (*aimP*) and receive (*aimR*) a signal peptide. AimP is secreted and cleaved extracellularly into the mature arbitrium signal peptide, typically consisting of 6 amino acids, which is internalised through the host-expressed OPP channel ^5^. AimR acts as an antiterminator ^9^ and binds to the upstream region of *aimX* in the absence of arbitrium signal, promoting the expression of *aimX* which promotes lysis; but as the concentration of signal peptide increases, AimR binds instead to the signal peptide, which inhibits expression of *aimX* and promotes lysogeny. Levels of *aimX* transcription control lysis/lysogeny via modulation of the host encoded MazEF toxin-antitoxin system ^9,12,13^. MazF activity promotes lysogeny likely by cleaving transcripts involved in the lytic cycle ^12^.

Theoretical and experimental studies have helped to explain how the arbitrium communication system can help phage to optimise their lysis/lysogeny decision, as the concentration of arbitrium signal correlates with susceptible host availability, and hence selection for lysis or lysogeny: AimP is produced during lytic replication and increases in concentration over the course of a phage epidemic, correlating with the decline of susceptible hosts for phage progeny to infect ^5,14^. AimP is also produced during the lysogenic cycle and broken down by proteases produced by host cells in a density dependent manner, thus an influx of susceptible hosts reduces the peptide concentration and prompts lysis ^6,7,10^. Obtaining information about the availability of uninfected hosts in the environment provides a key fitness advantage to arbitrium-phages ^6,14^. However, these studies have only considered the simple scenario where a clonal population of bacteria is infected by a single phage. If and how the fitness consequences of viral communication depend on the presence of other phages, and how the coexistence of different phages in the environment potentially shapes the evolution of phage communication systems, remains unclear.

Since the original discovery of the arbitrium systems, it has become clear that these systems are widespread in mobile genetic elements (MGEs) that infect *Bacilli* and other Firmicutes ^15^. Interestingly, these MGEs encode a large diversity of *aimR* receptors that have been grouped into nine clades based on their genetic relatedness, each associated with a distinct set of signal peptides that are predicted to range from 6 to 10 amino acids in length, depending on the clade; clade 2 arbitrium systems include Phi3T and SpBeta phages and have a 6 amino acid signal peptide ^15^. Experimental studies suggested that arbitrium systems are highly specific, as phages SpBeta and Phi3T carrying arbitrium systems belonging to the same clade only responded to their own cognate signal ^5^. The N-terminal region of the clade 2 peptides vary, but the C-terminal region is often conserved (RGA) i.e. SAIRGA for Phi3T and GMPRGA for SpBeta. Resolved crystal structures of signal-bound AimR from Phi3T and SpBeta have identified the receptor residues that bind to the signal peptide and shown that the amino acids that interact with the RGA motif of the signal are highly conserved across the clade 2 AimR receptors^8,16^.

However, the current consensus that arbitrium systems only respond to their own signal is based on a very limited number of examples, and more systematic studies are required to understand whether this is a general feature of these communication systems. We hypothesized that controlling the lysis/lysogeny switch with signal peptides may carry a risk of behavioural manipulation by other phages in the environment. To explore this, we first established that arbitrium-carrying phages naturally exist as polylysogens within the same host genomes, indicating that interactions between different phages with arbitrium communication systems are likely to occur in nature. Moreover, we found that phages may carry similar signals even if they belong to different species or genera, due to the frequent horizontal transfer of these communication systems. Next, we analysed the degree of cross talk within and between arbitrium clades, using the clade 2 systems as our model which include the well-studied phages SPbeta and Phi3T that infect *Bacillus subtilis*. Our experiments show that phages commonly influence each other’s lysis/lysogeny decisions through cross talk, leading to early lysogeny of the signal receiving phage, and elevated fitness of the signal-emitting phage. Our data suggest that antagonistic coevolution between signal emitting and signal receiving phages may drive the evolution of diverse and unique signals and receptors.

## Results

### Co-occurrence of Bacillus phages that encode arbitrium systems

To explore the potential for cross talk we first wanted to know if arbitrium-carrying *Bacillus* phages are likely to be exposed to each other’s signals in nature. We considered that if two arbitrium-carrying phages formed polylysogens then this would provide good evidence that they will be exposed to each other’s signals. By re-analysing the supplementary data from ((Stokar-Avihail et al., 2019) (their table S1)) containing homologs of the Phi3T *aimR* gene, their prophage and host, filtered for high quality phage assemblies only (N = 546 *aimR* sequences from 328 unique genomes), we found that while more than half of the bacterial genomes (54.3% (178/328 unique genomes)) carried only a single arbitrium-carrying prophage, polylysogeny was common with 35% (116 of 328) carrying two such prophages. 4.3% and 3.9% of genomes carried three and four arbitrium prophages, respectively, while a single genome contained eight prophages (figure S1). Of those genomes that carried more than one arbitrium prophage, there were clear species-specific patterns in the combination of arbitrium systems that polylysogens carried (figure 1a, table S1), with phages found in the *B. cereus* species group (consisting of *B. anthracis, B. cereus, B. thuringiensis*) generally carrying combinations of clades 4, 6 and 9 arbitrium systems, whereas those in *B. subtilis* carried clade 1, 2 and 3 systems. Furthermore, 43 genomes carry multiple prophages with arbitrium systems belonging to the same clade; 63% of these 43 are found in prophages of *B. thuringiensis,* a species that commonly harbours multiple arbitrium phages (figure 1a). Overall, these data show that in their natural environments, phages that use arbitrium signalling to control their lysis/lysogeny switch will not only be exposed to their own signals, but also to those of other phages, and these signals can belong to the same or other clades of arbitrium systems.

**Figure 1.**
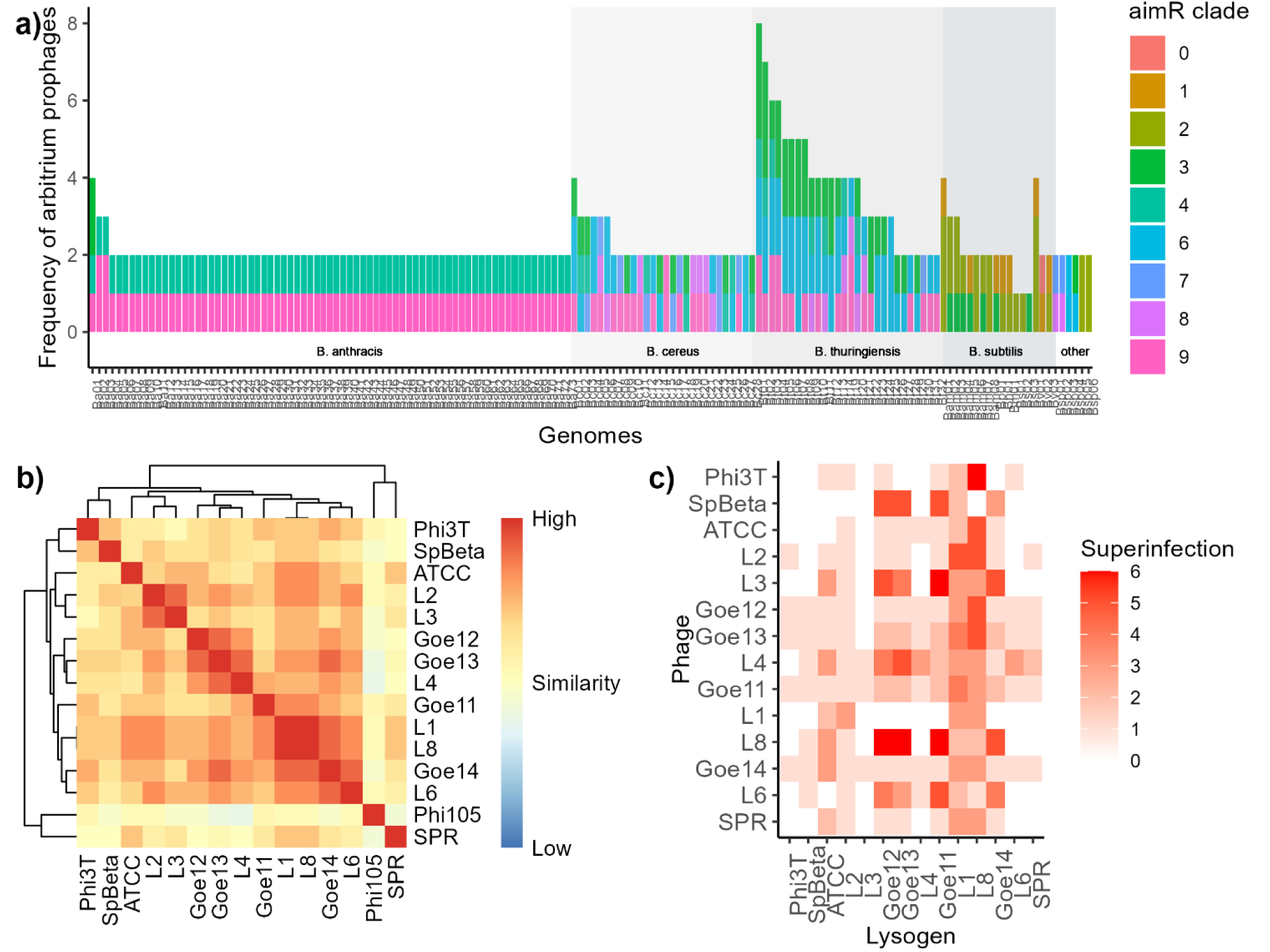
Arbitrium phages encounter each other in nature a) Bacterial genomes harbouring more than one arbitrium carrying prophage coloured by *aimR* clade (IMG genome IDs in table S1), b) Host range (Jaccard) similarity of a collection of 14 clade 2 SpBeta phages and Phi105 (no arbitrium) based on infection dynamics growth curve data of each phage against a panel of 30 *B. subtilis* hosts (figure S2 for raw data) c) Superinfection ability of 14 clade 2 phages against *B. subtilis* 168Δ6 lysogenised by the same phages.

To experimentally test whether natural phages carrying arbitrium systems belonging to the same clade can co-infect the same hosts, we analysed the host range of 15 *B. subtilis* phages that all carry clade 2 arbitrium systems. This revealed that many of the phages had a similar host range (figure 1b, figure S2), despite often low intergenomic similarity (ranging from 33.4 – 86.5% ani, figure S3). Further, we found that although some prophages provided protection from superinfection by other phages, all lysogens were susceptible to infection by at least some SpBeta phages (figure 1c). This, along with our polylysogeny analysis, shows that phages that carry arbitrium systems will frequently interact with both closely and distantly related phages that carry arbitrium systems of the same clade.

The co-occurrence of different arbitrium systems in the same bacteria through polylysogeny, shared host range and superinfection provides the opportunity for horizontal gene transfer (HGT) of systems between phages. A phylogenetic comparison of the *aimR* gene from each phage genome against the conserved *large terminase* gene revealed that the two trees are topologically very different i.e. the taxa group together in very different ways depending on which gene (*aimR* vs *large terminase*) was used to build the tree (Robinson-Fould distance = 0.9) (figure S4) ^17^, which suggests frequent horizontal acquisition of arbitrium systems, despite sharing weak similarity in branch distances (Mantel test, r = 0.21, p <0.01) ^18^. This helps to explain how even distantly related phage species that share an overlapping host range can harbour similar arbitrium systems (figure S3), and why we see bacterial species-specific arbitrium clade combinations (figure 1a).

### Phages can respond to non-cognate signals

To understand whether the lysis/lysogeny switch of Arbitrium phages are affected by non-cognate signals, we exposed *B. subtillis* strain 168Δ6 to a signal-deficient mutant of the model phage Phi3TΔ*aimP* in the presence or absence of a panel of 30 synthetic signal peptides from across eight arbitrium clades. This revealed that Phi3TΔ*aimP* significantly responds to its native SAIRGA plus four non-cognate signals - SIIRGA, SASRGA, SPSRGA and GVVRGA (figure 2). These four signal peptides are all produced by clade two arbitrium systems and are closely related to Phi3T’s cognate SAIRGA signal. Eight-hour growth curves following the infection dynamics of Phi3TΔ*aimP* and its host *B. subtillis* 168Δ6 show that these four non-cognate signals significantly reduce virulence of Phi3T infection, while the remaining 11 clade 2 signals, and 14 signals from other clades, have limited impact on infection dynamics (figure 2a, table S2 for Pairwise Wilcoxan Rank-Sum analysis).

**Figure 2.**
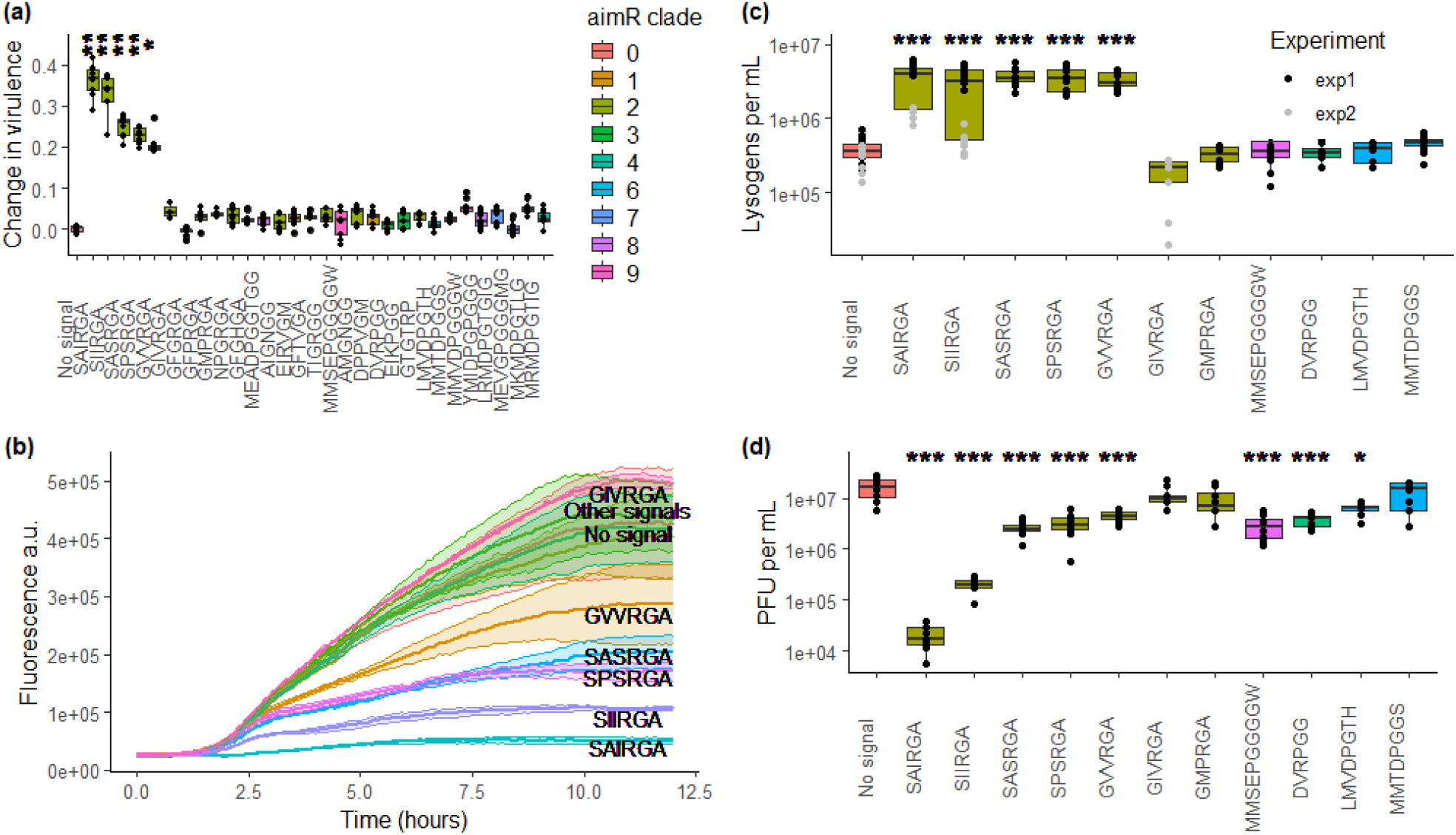
The Phi3T arbitrium system cross talks with non-cognate signals a) The effect of synthetic peptides on infection dynamics of Phi3TΔ*aimP* and *B. subtilis* 168Δ6; b) Fluorescence levels of a reporter strain that expresses *aimR* and *gfp* under the control of the *aimX* promoter in the presence of synthetic peptides, normalised to media only controls; c) The number of lysogens formed during the first round of infection (30 minutes) of Phi3TΔ*aimP*.spec in *B. subtilis* 168Δ6 in the presence of synthetic signals d) Prophage induction from *B. subtilis* 168Δ6::Phi3TΔ*aimP* lysogens in the presence of synthetic signals. Asterisks indicate the statistical significance of each treatment compared to the no-signal control, Pairwise Wilcoxon rank-sum test adjusted p-value: * <0.05, **<0.01, ***<0.01.

To demonstrate that this effect was mediated through a direct interaction between the non-cognate signals and the AimR receptor, we used a reporter assay where GFP was fused to the Phi3T *aimX* gene to assess the Phi3T AimR response to each signal outside of the context of a phage infection: Decreased fluorescence indicates that the peptide binds to AimR. This assay demonstrated that the Phi3T receptor responds to the same four non-cognate signals (SIIRGA, SASRGA, SPSRGA and GVVRGA), whereas the other signals were comparable to no signal control and did not affect fluorescence (figure 2b, table S3 for Pairwise Wilcoxan rank-sum analysis). The effect of these four non-cognate signals on the expression levels of the reporter gene varies in a manner that is consistent with their effects on the infection dynamics (figure 2a and b). As expected, signal mutations (relative to the cognate SAIRGA signal) with a pronounced impact on either the polarity of amino acids or the length of their side chains of the first three residues of the peptide were generally associated with a more significant reduction in cross talk with the Phi3T arbitrium system (table S4).

Next, we wanted to explore in more detail how non-cognate signals impact lysis/lysogeny decisions during different stages of the phage lifecycle. To this end, we used a smaller panel of synthetic signals (seven signals from clade 2 and four signals from clades 4, 8 and 9) to test their effect on Phi3T’s decision to enter or exit lysogeny. We found that SAIRGA and the same four non-cognate signals caused a significantly higher number of lysogens during the first round of infection, compared to a no signal control and other signals (figure 2c, table S5). Moreover, the presence of native SAIRGA and non-cognate SIIRGA peptides caused a significant decrease in prophage excision, compared to a no signal control (figure 2d, table S6), whereas SASRGA, SPSRGA and GVVRGA had a weaker (although statistically significant) effect on prophage excision, consistent with their weaker effect in the reporter assay. Several unrelated peptides that had shown no effect in other assays also weakly impacted prophage excision, suggesting this is not mediated through a direct interaction with the Phi3T AimR receptor.

### Cross talk is widespread and can be unidirectional or bidirectional

To investigate whether other arbitrium systems also cross talk, we first engineered Phi3T to encode the *aimR* and *aimP* genes and the *aimX* promoter from two other arbitrium systems that employ the SIIRGA and SASRGA signals, instead of the native arbitrium system. As before, we analysed the degree of cross talk using OD600 growth curves in the presence or absence of synthetic signals and calculated virulence over time from the area under the growth curves. As expected, this revealed that the Phi3T.SIIRGA mutant responds most strongly to SIIRGA, instead of the Phi3T cognate signal SAIRGA, and the Phi3T.SASRGA mutant responds mostly strongly to SASRGA. Crucially, the allelic replacement of the arbitrium systems showed that the existence of cross talk is not unique to the arbitrium system of Phi3T but the specific patterns of cross talk differ between the different arbitrium systems (figure 3a).

**Figure 3.**
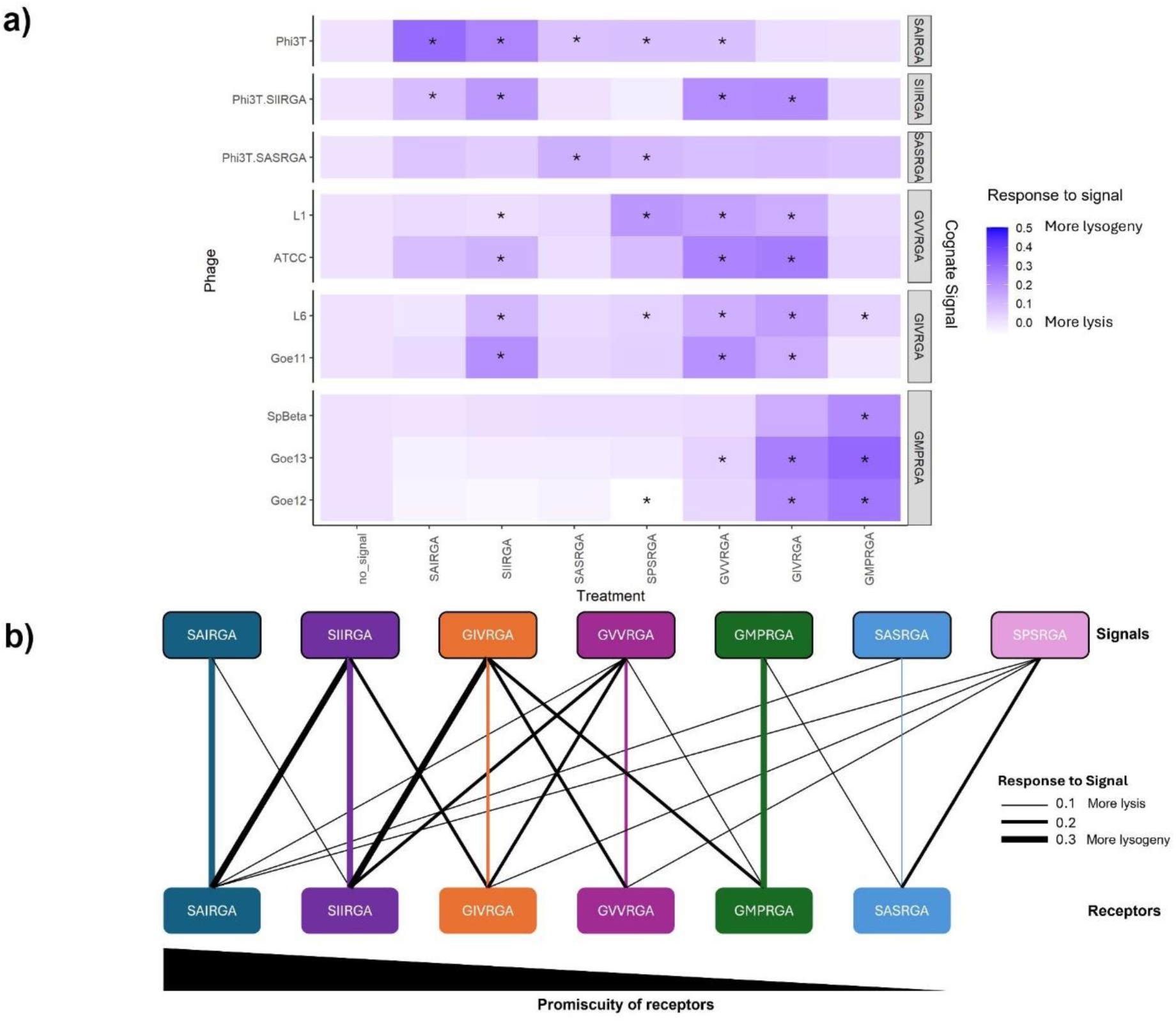
Cross talk is a general feature of arbitrium communication systems a) The effect of synthetic peptides on infection dynamics of a panel of natural and mutant phages and *B. subtilis* 168Δ6, measured as virulence levels relative to a no signal control; asterisks display significance of the treatment compared to the no signal control, Tukey-adjusted p value < 0.05, b) Schematic network of interactions between receptors and signals, ordered by receptor promiscuity to signals, based on the virulence data. Coloured arrows indicate interactions with self, whereas black arrows indicate crosstalk. The weight of each arrow represents the strength of response of each receptor to each signal.

To further generalise our finding that arbitrium systems respond to non-cognate signals, we also analysed cross talk in natural phages beyond our Phi3T model. Specifically, we infected *B. subtillis* strain 168Δ6 with seven natural SpBeta-like phages that are genetically diverse (figure S3) and that carry three different clade 2 arbitrium systems. This further reinforced our finding that phages with arbitrium systems commonly respond to non-cognate peptides (figure 3a,b, table S7) and revealed that arbitrium systems vary in their promiscuity within this set of peptides (figure 3b). Crucially, phages that carry arbitrium systems that employ the same signal peptides also show similar patterns of cross talk (figure 3a), even if the phages themselves have overall limited sequence similarity (between 47.1 – 86.5% ani, figure S3). Intriguingly, cross talk can be either unidirectional or bidirectional; for example all GMPRGA systems respond strongly to GIVRGA peptide, but GIVRGA systems respond weakly, if at all, to GMPRGA peptides (figure 3a).

### Manipulation of the lysis/lysogeny switch when phages infect different cells

Having established that arbitrium phages can respond to a broad range of non-cognate signals, we next wanted to understand if and how arbitrium signals when produced at ecologically relevant concentrations during a phage epidemic shape the lysis/lysogeny decisions of other phages. To test this, we infected cultures of *B. subtilis* 168Δ6 bacteria with either Phi3TΔ*aimP* (preferentially lytic) or Phi3TΔ*aimRPX* (preferentially lysogenic) mutants (both producing no signal, serving as negative controls), WT Phi3T (producing SAIRGA), Phi3T.SIIRGA (producing SIIRGA), phage Goe11 or Goe14 (both producing GIVRGA), and Goe12 or Goe13 (both producing GMPRGA). We then filtered the media to remove cells and phage, and used these “conditioned media” to grow and infect cultures of *B. subtilis* 168Δ6 with either WT Phi3T phage (producing and responding to the SAIRGA signal) or the isogenic Phi3T.SIIRGA mutant (producing and responding to the SIIRGA signal). Growth curve data confirmed that both phages had significantly reduced virulence in media conditioned by their cognate phage (WT Phi3T and Phi3T.SIIRGA, positive controls), and neither phage responded to media conditioned by the no signal negative controls (Phi3TΔ*aimP* and Phi3TΔ*aimRPX*) or the media conditioned by the GMPRGA producing phage (Goe12 and Goe13). However, Phi3T.SIIRGA displayed reduced virulence in response to conditioned media from GIVRGA producing phages Goe11 and Goe14 phages (figure 4a, table S8, figure S5), whereas WT Phi3T did not show a response to this signal, consistent with the cross talk patterns obtained with synthetic signals (figure 2 and 3). These data show that signal produced by one arbitrium phage can alter the infection dynamics of other phages in the same environment, leading to an increase in lysogeny of the cross talking phages.

**Figure 4.**
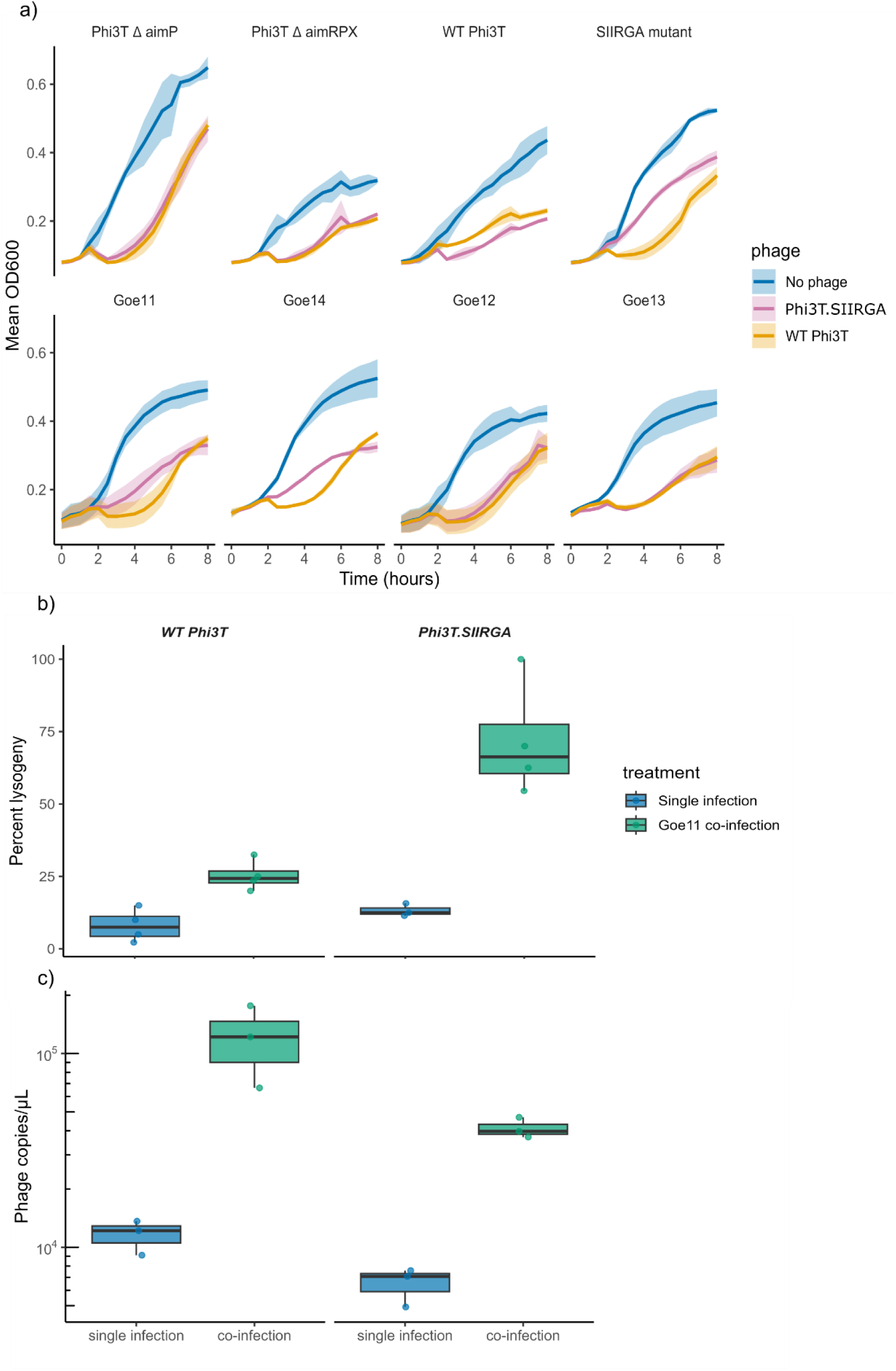
Cross talking phages influence each other’s infection dynamics a) Infection dynamics of WT Phi3T versus the Phi3T.SIIRGA mutant in media conditioned by phages carrying different arbitrium systems - no signal from Phi3T Δ*aimP* and Δ*aimRPX* (negative controls), SAIRGA from WT Phi3T (positive control for WT Phi3T infection), SIIRGA from Phi3T.SIIRGA mutant (positive control for Phi3T.SIIRGA infection), GIVRGA from Goe11 and Goe14, or GMPRGA from Goe12 and Goe13 (negative controls) b) Free phage single infections (WT Phi3T, Phi3T.SIIRGA) and co-infections (WT Phi3T + Goe11 or Phi3T.SIIRGA + Goe11) of *B. subtilis* 168Δ6 host, measuring percentage lysogeny and c) phage copy number in each treatment at 2 hours post-infection.

As the conditioned media experiments demonstrated that signal produced by past phage epidemics influence subsequent phage infection dynamics, we then wanted to understand if similar effects would occur when different phage simultaneously infect a bacterial population. To this end, we infected populations of *B. subtillis* 168Δ6 with both Goe11 (GIVRGA producing phage) and either WT Phi3T or Phi3T.SIIRGA as free phages. This revealed that many more lysogens were being formed by Phi3T.SIIRGA at two hours post infection (mean 71% lysogeny) compared to the WT Phi3T (mean 25% lysogeny) (Pairwise Wilcoxan rank-sum test, W = 0, p = 0.029, Bonferroni p.adj = 0.058, effect size = 0.8) during co-infection with Goe11, with the WT Phi3T associated with a non-significant increase in the production of free Phi3T phage particles (W = 9, p.adj = 0.2) (figure 4b). As expected, there was no difference in lysogeny formation between the phages in the *B. subtilis* 168Δ6 background (negative control) (figure 4c) (W = 2, p = 0.2, p.adj = 0.5, effect size = 0.5). Polylysogeny was rare under these experimental conditions (figure S6), which suggests that a signal released by one phage can be detected by others infecting different cells, leading to increased lysogeny when there is cross talk between arbitrium systems.

### Manipulation of the lysis/lysogeny switch when phages co-infect the same bacterial cell

As many *Bacillus* genomes contain at least one arbitrium prophage (figure 1a), and lysogens are known to produce arbitrium signal ^6,7,10^, we hypothesised that cross talking signals produced by a lysogen could also manipulate the lysis/lysogeny switch of a superinfecting phage. To test this, we lysogenised *B. subtilis* 168Δ6 with Goe11 (GIVRGA producing phage). First, we confirmed that Goe11 lysogens were susceptible to infection by Phi3T (figure 1c). Next, we used these lysogens and WT *B. subtilis* 168Δ6 in infection experiments with either WT Phi3T or the Phi3T.SIIRGA mutant. We found that the efficiency of polylysogeny in the *B. subtilis* 168Δ6::Goe11 host after the first round of infection (30 minutes) was 10-fold higher for Phi3T.SIIRGA, compared to the isogenic WT Phi3T (Pairwise Wilcoxan rank-sum, W = 81, adj. p-value < 0.001, effect size = 0.84), while there was no difference between the phages in percentage lysogeny of the WT *B. subtilis* 168Δ6 host (W = 62, adj. p value = 0.8, effect size = 0.2) (figure 5a). These data show that the GIVRGA signal produced by the *B. subtilis*Δ6::Goe11 lysogen enhances (poly)lysogeny of the cross talking Phi3T.SIIRGA phage, but not the WT Phi3T - consistent with our findings using synthetic signals, conditioned media and free phages. Crucially, while early lysogeny can reduce fitness for the receiving free phage by limiting its capacity to lyse and replicate, it benefits the signal-emitting prophage by encouraging lysogeny and deflecting lytic infection (figure 5b).

**Figure 5.**
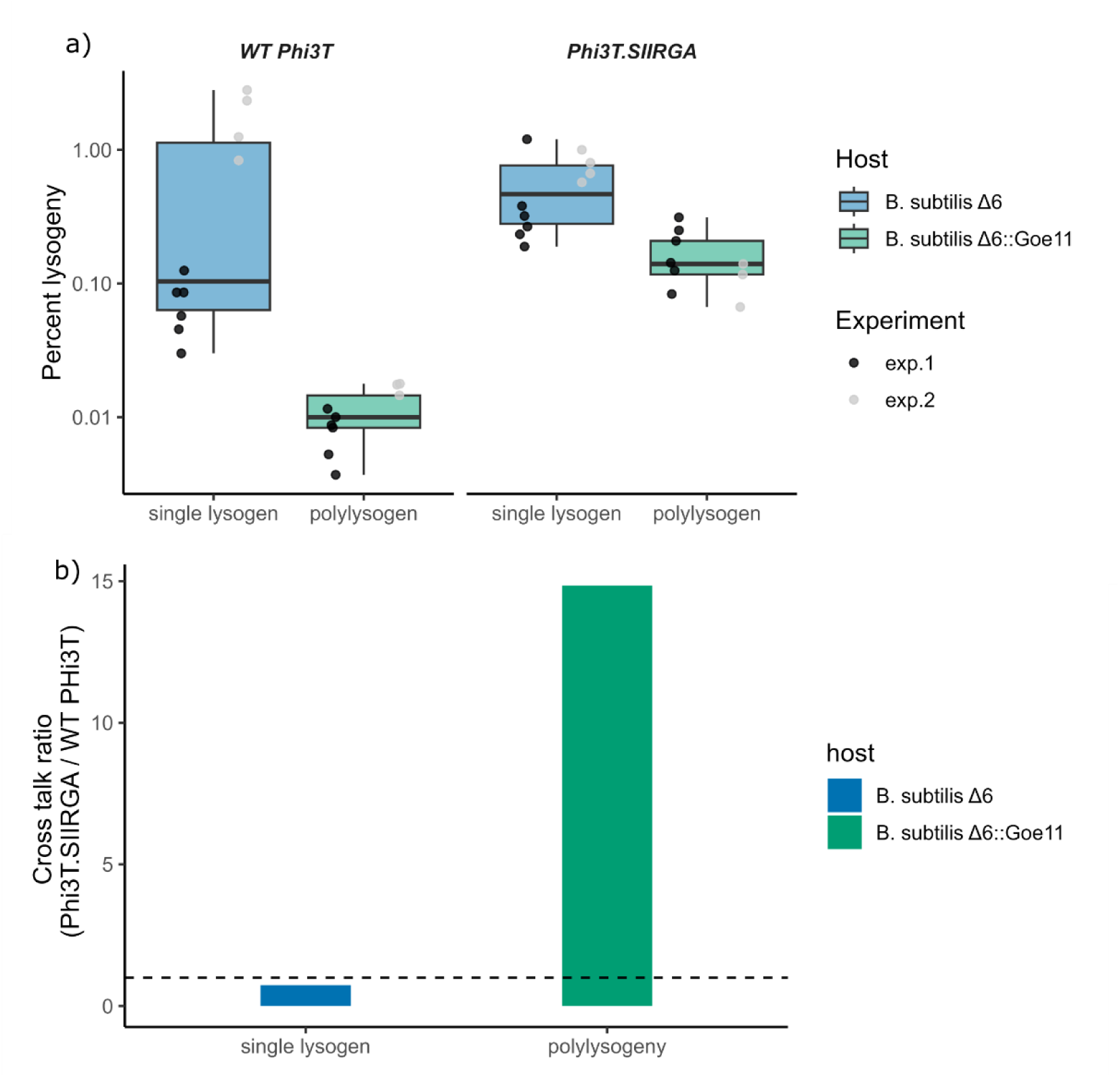
Cross talk with a prophage can alter infection dynamics causing the receiving phage to lysogenise early, but with fitness benefits to the signal emitting prophage a) Efficiency of lysogeny of WT Phi3T and Phi3T.SIIRGA mutant after the first round of infection (30 minutes) in the prophage free strain *B. subtilis* 168Δ6 and *B. subtilis* 168Δ6::Goe11 lysogen hosts, b) Cross talk ratio, calculated as the mean polylysogeny of *Phi3T.SIIRGA* divided by that of wild-type *Phi3T* in both *B. subtilis* 168Δ6 and *B. subtilis* 168Δ6::Goe11 backgrounds. The dotted line at ratio = 1 reflects equal lysogeny rates between phages, and ratios > 1 indicate that the signal-emitting prophage manipulates the superinfecting phage to lysogenise the host.

## Discussion

Our findings show that naturally occurring phage-derived arbitrium peptides can manipulate the lysis-lysogeny switch of distantly related phages that carry similar arbitrium systems, promoting early lysogeny and reducing prophage induction. In recent years, the concept of sociovirology, where viruses display “social” behaviours to enhance their fitness, has received a lot of attention^19,20^. Our finding that viruses can manipulate each other’s infection dynamics highlights potential risks associated with viral cooperative behaviours. Cross talk will generally be maladaptive for the signal receiving phage ^21^, as it causes the phage to underestimate the density of available hosts in the environment and initiate lysogeny prematurely. Negative fitness consequences of cross talk will favour the invasion of mutants that are resistant to cross talk through the evolution of a novel signal specificity, which is consistent with the extensive diversity of *aimR* genes found across *Bacillus* phages^15^ and horizontal exchange of arbitrium systems between phages, indicating evolution away from cross talk. The observed cross talk may therefore be a transient state towards the evolution of unique signal-receptor pairs.

Future studies are needed to shed light on the mutational pathways that allow for the evolution of novel specificities. Arbitrium systems share some key similarities with *Bacillus* Rap-Phr quorum sensing systems, which also belong to the RRNPP family of quorum sensing systems ^22^ and these are also highly diverse. In the case of these bacterial quorum sensing systems, diversity is thought to result from *phr* duplication and mutation of the duplicated *phr*, followed by co-evolution of the *rap* receptor gene to generate a novel Rap-Phr specificity and loss of the ancestral *phr* ^23^. Possibly, arbitrium systems diversify through similar mechanistic steps. Premature lysogeny would create a selection pressure for mutants to produce a cognate signal with a lower affinity for *aimR*, supporting more lytic replication.

As the concentration of the novel signal peptide builds up, lysogeny becomes optimal, and the co-evolution of *aimR* would result in a shift in specificity. Alternatively, the *aimR* gene may mutate first to reduce the affinity for the non-cognate signal, and the *aimP* gene may coevolve with the mutated *aimR*. It only takes a single amino acid substitution in AimR to alter its specificity for the AimP signal peptide ^8^, a feature also observed in another RRNPP family member, RapF ^24^, which implies arbitrium systems can shift signal peptide affinity through minimal genetic changes to the *aimR*. That said, there are likely mechanistic constraints that prevent the evolution of perfect specificity in these signal-receptor interactions ^25^, which, along with the stepwise nature of the coevolution between the signal and receptor, may explain why cross talk is observed at high frequencies in nature.

While cross talk undermines signal fidelity and will often be costly for the signal-receiving phage, it can provide a clear fitness advantage to the signal-emitting prophage. Indeed, if a prophage encourages a superinfecting phage to enter into lysogeny, rather than lysing the bacterial host, it directly reduces the mortality risk for the signal-emitting phage. Whereas manipulation of host behaviour by viruses and other parasites is well-documented ^26^, the observation that viruses can use signals to influence each other’s behaviours to enhance their transmission provides a novel twist on the factors that shape viral epidemiology. Furthermore, the evolutionary downsides of using communication systems to regulate lysis– lysogeny decisions in a community context may counterbalance some of the fitness advantages these systems provide, such as improved decision-making based on fluctuating densities of uninfected bacteria ^6^. Understanding the fitness benefits of viral communication, but also that these systems can come at a cost, may help to guide future efforts to identify analogous systems elsewhere.

## Methods

### 1. Experimental analyses

#### Bacterial strains and culture conditions

All bacterial strains used in this study belong to *B. subtilis* (table S9) or *E. coli* species. All strains were routinely grown at 37C on Luria Broth (LB) agar plates or in LB liquid medium, with shaking at 200 rpm. When required, antibiotics were added at the following concentrations unless stated otherwise: kanamycin (5 µg/mL for *B. subtilis* or 30 µg/mL for *E. coli*), ampicillin (100 µg/mL), tetracycline (10 µg/mL) or spectinomycin (100 µg/mL). *B. subtilis* 168::Δ6, a strain of *B. subtilis* 168 with two prophages (SpBeta, PBSX), three prophage-like regions, and the largest operon of *B. subtilis* (pks) genetic elements deleted ^27^ was kindly provided by Prof. Jose Penades and used throughout this work to eliminate the possibility of interference by these MGEs.

Luria-Bertani (LB) agar and broth was used for bacterial growth unless otherwise stated. LB broth and agar were supplemented with 0.1mM MnCl2, 5mM MgCl2 MD for phage infections. All cross talk experiments were carried out using MD media: 10x PC was prepared and autoclaved: 0.6M K_2_HPO_4_, 0.4M KH_2_PO_4_ and 19mM trisodium citrate dihydrate Na₃C₆H₅O₇·2H₂O), then MD media was prepared to final concentration of 1xPC, 0.11M glucose, 0.25mM L-tryptophan, 14.6mM aspartic acid potassium salt, 3mM MgSO_4_, 0.1% casamino acids and freshly prepared 37.5µM ferric ammonium citrate). M9 x1 was used as a phage buffer for dilutions: 5x stock - 42mM Na_2_HPO_4_, 22mM KH_2_PO_4_, 9mM NaCl and 19mM NH_4_Cl.

#### Natural phages and amplification

Phages Phi3T, SpBeta and Phi105 were purchased from the Bacillus Genetic Stock Centre (BGSC), SpBeta.L1, L2, L3, L4, L6 and L8 were a gift from the Eldar and Sorek labs, Goe11, Goe12, Goe13 and Goe14 were gifted from the Penades lab. High titre phage stocks were obtained from lysogens and stored at 4C: lysogens were allowed to grow to 0.2 OD600, before being treated with 5µg/mL Mitomycin C and kept at 30C at 80rpm for 4 hours. After an overnight at room temperature overnight cultures were filtered through 0.22 µM pore syringe filters (PES membrane, Merck, Millipore) to remove bacteria. Titres were measured using traditional top agar overlay with *B. subtilis* 168Δ6 and spotting on phage dilutions.

#### Signal peptides

30 signal peptides were synthetically made by Peptide 2.0, ISCA Biochemicals Ltd or GenScript and hydrated to 1mM concentration stocks (table S10). The signals were chosen based on the clades of the *aimR* phylogeny by ^15^. 16 signals were chosen from clade two (related to Phi3T); and two were chosen at random from each of the other seven clades. Signals were used at a final concentration of 1000nM in all assays.

#### Construction of reporter plasmid

The plasmid DH5alpha::pDR111_*aimRPX*_sfGFP, a gift from Gil Amitai of Rotem Sorek’s lab, containing the arbitrium genes from Phi3T (*aimR*, *aimP* and *aimX*), was modified in this study to remove the *aimP* gene, and transformed into *B. subtilis*::Δ6 where it was inserted within the *amyE* gene (table 9). *B. subtilis* Δ6::*aimRX*_sfGFP was subsequently used as a reporter strain to measure the response of Phi3T *aimR* to non-cognate signals. Briefly, bacteria carrying the plasmid were grown overnight in the presence of ampicillin (100ng/µL) and the plasmid was extracted using the Monarch Miniprep kit, following manufacturer’s instructions. The vector was double digested using EcoR1 HF and Bam HF (NEB) following manufacturer’s instructions, run on a 1% agarose gel and the 6000bp fragment was purified via freezing (-80C for 5 minutes) and centrifugal spinning through a 200µL filter tip, followed by ethanol precipitation. Using the extracted vector as a template, fragments upstream and downstream of the *aimP* gene were amplified by PCR using NEB Q5 polymerase (table S11), and single bands were cut out and DNA extracted as described above. The NEB HIFI assembly kit was used to re-assemble the vector without the *aimP* gene before transformation into DH5 alpha cells (NEB) (42C for 30 seconds) and plating onto ampicillin plates. The new vector was extracted as above, quantified by Qubit DS DNA broadrange assay, and transformed into competent *B.subtilis168*::Δ6. Briefly, 10 µL of overnight *B.subtilis*::Δ6 cells were diluted into 1mL MD media, grown for 3 hours at 37C and 200rpm; then 200ng of vector DNA was added to 300 µL of culture and left for another 3 hours incubation before plating onto kanamycin (5 µg/mL) plates. Individual colonies were picked and grown O/N in LB before storing the reporter strain in cryostocks.

#### Construction of phage mutants

Pjoe8999, a CRISPR-Cas9 *B. subtilis-E. coli* shuttle vector ^28^, which contains a single guide (sg) RNA sequence and *cas9* under the control of a mannose-inducible promoter, was used to engineer all changes to prophages within *B. subtilis* 168::Δ6 (table S12). Annealed and dephosphorylated sgRNA oligos targeting the region for deletion or insertion were first ligated into the Bsal HF (NEB) digested vector using T4 ligase (NEB). Arbitrium system mutants Phi3T::SIIRGA and Phi3T::SASRGA were constructed using sgRNA targeting the WT Phi3T *aimR;* the insertion sequences carrying the *aimRP-aimX-promoter-region* of the alternative (SIIRGA and SASRGA) signalling systems were synthesised as gBlocks (IDT) and inserted in the Sfil digested vector containing the sgRNA using the NEB HiFi DNA Assembly Cloning Kit (table S13). Vectors were transformed into competent *E. coli* DH5alpha cells, grown overnight on Kanamycin plates (50 µg/mL). Colonies were selected and vectors extracted using a miniprep kit (Monarch) and transformed into *B. subtilis* 168::Δ6 carrying WT Phi3T as a lysogen. Colonies were screened using antibiotics and heat treatment steps as described in Altenbuchner 2016. Briefly colonies were first grown at 30C for 2 days, then 50C overnight and finally 42C overnight before checking for plasmid loss on kanamycin. Recombination was confirmed by whole genome sequencing of the lysogen (Microbes NG).

Kanamycin or spectinomycin resistance cassettes were introduced into the Phi3T genome, replacing the non-essential Phi3T_5 (*yokI*) ^10^. 1000bp regions flanking Phi3T_5 and either a kanamycin or spectinomycin resistance cassette were amplified and introduced into the sfiI-digested vector using NEBuilder HiFi DNA Assembly Master Mix (table S13). The resulting constructs were transformed into *E.coli* DH5alpha cells (NEB) for amplification before transformation into *B*. *subtilis* 168Δ6 harbouring the Phi3T prophage to generate the desired mutants. Individual colonies were screened using PCR to identify cells containing the antibiotic resistance cassette. Lysogens were then reactivated, and individual phage mutants plaques picked for further testing. Phage infections were carried out, and cultures plated onto LB agar containing either kanamycin or spectinomycin to confirm that the phage antibiotic resistance cassette was conferring antibiotic resistance to the susceptible host. The Goe11 mutant containing a tetracycline cassette also replacing yokI, Goe11::Δ*yokI*::*tetL*, was a gift from the Penades lab.

#### Host range and superinfection analysis

To assess the host range of each phage against a panel of 30 hosts (27 *B. subtilis* and 3 *B thuringiensis*) (table S14), each host was grown up overnight (O/N) from a single colony, diluted 1:100 and grown at 37C, 200rpm, until OD600 = 0.2. ∼1x10^6^ cells were added to an Eppendorf tube and phages were added at an MOI of ∼ 0.1. Controls containing no-phage were run for each host, and media-only controls were run on every plate. Infection assays were run in single wells per host-phage pairwise comparison. Plates were incubated at 37 °C for eight hours with shaking before each optical density (OD_600_) measurement was taken every 30 minutes on a microplate reader (Infinite 200 Pro, Tecan). The virulence of phages against each host was calculated from the growth curves as 1 – area under the curve (AUC) phage treatment/AUC no phage treatment ^29^. A binary measure of infection was created using a cut off of virulence = 0.2 (< 0.2 = no infection, > 0.2 = infection). Jaccard similarity was computed based on host range profiles using the vegdist() function (method = "jaccard") from the *vegan* package in R (v. 2.7)^30^. Resulting dissimilarities were converted to similarity values (1 - distance) and visualized using a heatmap.

To assess superinfection of each phage against each other’s lysogens, first lysogens were made from each phage in *B. subtillis* 168Δ6 and each lysogen was tested by spotting onto *B. subtillis* 168Δ6 top agar and, after O/N incubation, observing lysis. Each lysogen was diluted into top agar and each phage spotted on in 1:10 dilution series, with one biological rep per host-phage pair. The plates were grown overnight at 37C. Plaques and areas of lysis were assessed based on a modified previously published superinfection scoring index ^31^:

0 - No spots of lysis or plaques
1 - Spots of lysis at highest 1–2 titres, but no plaques
2 - Superinfecting phage produces plaques with an efficiency of plating of less than 10^3^–10^4^, or spots of lysis at highest 3 titers but no plaques
3 - Superinfecting phage produces plaques with an efficiency of plating from 10^1^–10^3^ or spots of lysis at highest 4–5 titres but no plaques
4 - Superinfecting phage produces plaques with an efficiency of plating of 1, but spots/plaques exhibit increased turbidity or reduced size compared to infection of *B. subtilis* 168Δ6
5 - Superinfecting phage produces plaques with an efficiency of plating of 1, and there is no phenotypic difference compared to infection of *B.subtilis168Δ6*
6 - Superinfecting phage produces plaques with an efficiency of plating of 1, but spots/plaques exhibit reduced turbidity or increased size compared toinfection of *B. subtilis* 168Δ6.

#### OD_600_ growth curve assays

To measure the response of Phi3T and other phages (Goe11, Goe12, Goe13, L1, L6, ATCC, Phi3T, SpBeta, and the two arbitrium Phi3T mutants SIIRGA and SASRGA) to synthetic signals, MD media was inoculated with ∼1x10^6^ *B. subtilis* 168Δ6 cells (exponential phase, optical density (OD)_600_ ∼ 0.2) and ∼1x10^5^ plaque forming units (PFU) of phage to a final MOI of 0.1. Signal peptides (30 in the Phi3T assay and 7 in the other phage assays, were added as individual treatments at a concentration of 1000nM in triplicate to a 96 well plate to final volume of 200 µL per well. Controls containing media only, no signal peptide, and no phage, were run on each plate. Plates were incubated at 37 °C for eight hours with shaking before each OD_600_ measurement was taken every 30 minutes on a microplate reader (Infinite 200 Pro, Tecan). The virulence of phages against each host was calculated from the growth curves as 1 – area under the curve (AUC) phage treatment/AUC no phage treatment ^29^. The experiment was run with three biological replicates, the difference between virulence for each treatment and the no signal control was calculated per run (mean virulence of no signal control – virulence of treatment) before combining the data across runs.

#### Reporter Assay

An Eppendorf containing 600 µL of MD media was inoculated with ∼1x10^6^ *B. subtilis*::Δ6::*aimR*, *p_aimX_-gfp* (exponential phase, OD_600_ = 0.2) and 30 signal peptides and control treatments were run as above. Using a BMG Labtech CLARIOstar microplate reader, plates were incubated at 37 °C for eight hours with shaking before each measurement of OD_600_ and fluorescence (YFP: excitation was set to 514 nm, and emission was recorded at 527 nm) every 10 minutes for 12 hours. Data were normalised to the media only control.

#### Efficiency of lysogeny assays

*B. subtilis* 168Δ6 was grown in LB from a single colony overnight, before being diluted 1:100 and grown to OD_600_ ∼0.4 in four biological replicates. Cells were centrifuged at 4000 x g for 10 minutes, the supernatant removed, and cells resuspended in MD media to an OD_600_ of 0.2. 1.3mL of each culture was added to wells in a 24 well plate in 3 technical replicates, ∼ 5 x 10^7^ cells per well. 12 treatments included a no signal control and 11 signal peptides added at a concentration of 1000nM and cells were incubated for 30 minutes at 37C and 200rpm for signals to be absorbed by the cells, before infection with phage mutant Phi3TΔ*aimP*.spec at an MOI of 0.1 (∼5 x 10^6^ PFU). After 30 minutes, 1mL of culture per well was transferred to individual Eppendorfs, centrifuged at 7000 x g for 1 minute, the supernatant removed, cells washed in M9 x 1 and resuspended to remove free phages. Each sample was gently homogenised and diluted in a 1:10 dilution series and spotted onto LB plates containing spectinomycin (100ug/mL). Plates were incubated at 30C overnight and colonies were counted the following day.

#### Prophage induction assays

*B. subtilis168*Δ6::Phi3TΔ*aimP* lysogens were grown in LB from a single colony overnight, before being diluted 1:100 and grown to ∼0.4D_600_ in four biological replicates. Cells were washed as above and resuspended in LB to 0.3 OD_600_. 1mL of MD media was added to each well of a 24 well plate with 300µL of resuspended cells (∼6 x 10^6^ per well). Plates were incubated at 37C, shaking at 180rpm for 6 hours. After 6 hours, 200µL of culture was harvested from each well into a 96 well 0.45 µm pore filter plate (MERCK) and spun for 2 minutes at 900rpm. Each sample was diluted in a 1:10 series and spotted on to a top agar overlay of *B. subtilis* BEST7003:*aimX* with xylose added at 0.2%. Plates were incubated at 37C overnight and plaques were counted the following day.

#### Conditioned media experiments

Overnight culture of *B. subtilis168Δ6* was diluted 1:100 into 8mL MD media and allowed to grow at 37C, 200rpm for ∼3 hours until OD_600_ = 0.5, which equates to ∼ 1 x 10^8^ CFU/mL. Cultures were infected at an MOI of 1 with Phi3TΔ*aimP* and Phi3TΔ*aimRPX* (no signal control), WT Phi3T and Phi3T.SIIRGA (positive controls), Goe12 and Goe13 (GMPRGA, negative controls), and Goe11 and Goe14 (GIVRGA) as treatments. Infections were incubated for another 3 hours, then bacteria cells were centrifuged and filtered out. The resulting media, ‘conditioned’ by the previous phage infection, was used immediately in plate reader assays as described above (see “OD_600_ growth curve assays”). 1x10^6^ cells of fresh *B. subtilis* 168Δ6 host grown to OD_600_ 0.2 was added and infected with WT Phi3TWT.kan and Phi3T.SIIRGA.spc at an MOI of 0.1, or with no phage (control). Plates were incubated at 37C for eight hours with shaking before each OD_600_ measurement was taken every 30 minutes on a microplate reader (Infinite 200 Pro, Tecan). Samples were run across four plates, which were recorded and plotted as experiments 1-4. Virulence was calculated as above.

#### Co-infection experiments

To measure lysogen counts and free phage over time during co-infection of phages, an overnight culture of *B. subtilis168Δ6* was diluted 1:100 into LB media in 4 biological replicates and allowed to grow at 37C to OD_600_ ∼0.4. Cells were spun at 4000 x g for 10 minutes, supernatant removed, and cells resuspended in MD to an OD_600_ of 0.2. 1.3mL of each of the four cultures was added to wells in a 24 well plate, corresponding to ∼ 5 x 10^7 cells. Goe11 was added at a MOI =1 and either WT Phi3T.kan or Phi3T.SIIRGA.spc mutant were added as co-infections at an MOI of 0.1, alongside single infection controls. At 2 hours post infection, 1mL of each sample was pelleted and washed in M9 x1 to remove free phage, before being diluted in a 1:10 series and 5 µL plated onto selective plates with appropriate antibiotics to count lysogen and polylysogen formation, and to control for the absence of phage contamination. For free phage counts, at 2 hours post infection, 200 µL of each sample (N = 3) was filtered through a 96 well 0.45 µm pore filter plate and phage lysate was treated with DNase 1 (5mg/mL, Roche) and RNase A (10mg/mL, Invitrogen) for 30 minutes at 37°C to remove non-encapsulated (e.g. free bacterial DNA) nucleic acids from the lysate, enzyme activity was terminated with heat treatment ^32^ and dropletPCR was carried out using 2xQX200 Evagreen supermix and primers (table S15) targeting the kanamycin, spectinomycin cassettes of the Phi3T phages and an intergenic region after the aimX gene of the Goe11 phage. Primers were used at a final concentration of 100 nM and 2 µL of phage lysate was used diluted to 1:10 or 1:100 in a 20 µL reaction. No template controls were run on every plate. Droplets were generated using the QX200 Droplet generator before PCR amplification (Bio-rad): 5 minutes at 95C initial activation, and 40 cycles of 95C for 30s denaturation and 60C for 1min annealing and extension, followed by 4C for 5 min, 90C for 5min, before storing overnight in the dark at 4C. Fluorescence readings were measured the morning after on the QX200 Droplet Reader using Quantasoft software v.2.2 (Bio-rad laboratories). Only wells with at least 10,000 accepted droplets were included in the analysis. To verify assays a 1:10 dilution series of known titre was run for each primer pair.

#### Polylysogeny experiments

To measure the efficiency of lysogeny after the first round of infection of WT Phi3T.kan and Phi3T.SIIRGA.spc in *B. subtilis* 168Δ6 (control) and *B. subtilis* 168Δ6::Goe11.tet lysogen, the two strains were grown in LB from single colonies overnight, before being diluted 1::100 and grown to ∼0.4OD_600_ in 3 replicates per strain. Cells were spun at 4000 xg for 10 minutes, supernatant removed, and cells resuspended in MD to an OD_600_ of 0.2. 1.3mL of each of the three cultures was added to wells in three technical replicates in a 24 well plate ∼ 5 x 10^7^ cells. Each strain was infected with either WT Phi3T.kan or Phi3T.SIIRGA.spec at an MOI of 0.1 (∼5 x 10^6 PFU). At 30 minutes (p.i) 1mL of each sample was pelleted and washed in M9 x1 to remove free phage, before being diluted in a 1:10 series and 5µL plated onto LB plates and tetracycline for the *B. subtilis* 168Δ6 and *B. subtilis* 168Δ6::Goe11.tet assays, respectively, for total cell count; and on to both spectinomycin:tetracycline and kanamycin:tetracycline plates to count polylysogen formation, and to control for the absence of phage contamination. Plates were incubated at 30C overnight and lysogens counted the next day. This experiment was carried out twice. Efficiency of lysogen was calculated as the percentage of lysogens or polylysogens formed from the total cell count.

#### Statistical analyses

All statistical analyses were carried out in R (v.4.3.2) and R studio (v.2025.05.0) and plots were made with ggplot (v.3.3.5) ^33^. For OD_600_ growth curve assays of non-Phi3T phages, statistical analyses were conducted with an alpha level of 0.05. Data for each phage were analysed using linear mixed-effects models (with biological replicate included as a random effect). Non-significant fixed effects were removed sequentially using backward elimination. To determine the significance level of the treatment effect we compared models with and without the treatment term using likelihood ratio tests. Diagnostic plots (residuals vs. fitted, Q-Q, residuals vs. leverage and scale-location plots) were used to check model fit. Post-hoc pairwise comparisons were performed using the emmeans package ^34^, with significance determined using Tukey-adjusted p-values. Mean virulence values for each arbitrium system (across phages) in response to each signal were generated, and the promiscuity of each *receptor* was calculated from this receptor-signal interaction matrix using the species strength index from the Bipartite package ^42,43^. A schematic was then plotted to illustrate the network and strength of cross talk responses between arbitrium systems. For all other OD_600_ growth curve assays and the lysogeny and prophage induction assays, to test for statistical differences between the signal treatments compared to no signal, a Wilcoxon test from the package rstatix v. 0.7.0 ^35^ was used with p-values corrected for multiple comparison using the Bonferroni method, followed by the package coin v. 1.4.2 ^36^ calling wilcox_effsize, to calculate effect sizes and confidence intervals by bootstrap (nboot = 1000).

### 2. Bioinformatics analyses Polylysogeny analysis

To investigate polylysogeny of *aimR* carrying prophages within *Bacillus* genomes, we re-analysed the supplementary data available online (^15^ table S1) edited to remove gene contexts that were predicted as bacterial or unknown, and low quality phage genomes (i.e. incomplete genomes). Plots were made with ggplot (v.3.3.5) ^33^.

#### Phylogeny methods to investigate the evolutionary history of the *aimR* gene

To carry out a phylogenetic comparison between aimR and the conserved phage gene terminase L the phage genomes were obtained. Using the scaffold IDs provided in ^15^ (their table S1), full IMG genome IDs were obtained from the Integrated Microbial Genomes & Microbiomes (IMG/M) system at the DOE Joint Genome Institute (https://img.jgi.doe.gov/), and high quality prophage genomes from this bacterial dataset were pulled out, keeping only those genomes that were predicted to be high quality (> 50% complete) ^37^. From this viral dataset, prophages that contained an aimR matching the original table ^15^were extracted using blastp (v.2.15)^38^, with the original aimR sequences as a query file and a strict e-value cut off of 1e^-10^ resulting in 483 phage genomes. The genomes were annotated using Pharokka v. 1.3.0 (Bouras et al., 2023) with default settings ^40,41^. To look at the how related the phage genomes were vContact2 ^42^ was run on a concatenated file of the 483 phage proteins and clustering was visualised in Cytoscape ^43^ confirming that all the phages clustered together well enough to continue the analysis as a single cluster. *AimR* and *terL* genes from each phage were extracted from the Pharokka output files and aligned in Geneious10.1 with default settings. Phylogenetic trees were constructed using IQ-Tree v.2.2 ^44^ with the following parameters: ‘-msub viral -bb 1000 -mset’. Newick files were processed in R using ape v5.8 ^45^and plotted with ggtree v3.10.1^46^. Trees were pruned to aimR-*terL* matching IDs (N = 398). Congruence between *terL* and *aimR* phylogenies was tested in R using a Mantel test^18^ to assess similarity between the distance matrices of each tree, and a Robinson-Fould test ^17^to assess the difference in topology between the trees.

## Acknowledgements

We would like to thank Professor Sylvain Gandon and Dr Ryuichi Kumata from CNRS Montpellier for insightful discussions on the evolution of cross talk; and Dr Michelle Michelson and Dr Luis Bolanos of Exeter University for support in the lab and with phylogenetic analysis, respectively. We thank Dr Gil Amitai (Weizmann Institute, Israel) for kindly sharing with us the *aimRPX*-sfGFP reporter plasmid, and Prof. Penades and Prof. Eldar and their groups for sharing phages and strains with us.

## Author contributions

Conceptualisation, E.R.W, R.M and J.B.; Methodology, R.M, R.W, E.S, J.B, B.T, and E.R.W; Investigation, R.M, R.W. and J.B.; Formal analysis R.M, R.W, E.S. B.T.; Writing, R.M and E.W.; Review and editing, all authors; funding acquisition, E.R.W.

## Resource availability

### Lead author

Requests for further information and resources should be directed to and will be fulfilled by the lead contact, Robyn Manley.

### Materials Availability

Plasmids and phage mutants developed in this study are available by request to the lead contact.

### Data and code availability

- Raw data from all experiments will be deposited on Dryad and will be publicly available as of the date of publication at [DOI].
- This paper analyses existing, publicly available data from ^15^ in supplementary table 1, accessible at https://www.sciencedirect.com/science/article/pii/S1931312819301696#app2
- Sequence data of mutant phages and lysogens will be deposited on Genbank as [Database: accession number ###] and will be publicly available as of the date of publication.
- Any additional information required to reanalyse the data reported in this paper is available from the lead contact upon request.

## Supplementary tables and figures list

Figure S1: The number of arbitrium-carrying prophages within individual genomes

Figure S2: Virulence data of clade 2 phage panel against 30 *B. subtilis* hosts

Figure S3: Genomic comparison (Viridic) of the panel of clade 2 phage and their arbitrium systems

Figure S4: Comparison of phylogenetic trees of *aimR* and large terminase genes

Figure S5: Conditioned media data plotted as virulence calculated from the growth curves, which was subsequently used in the statistical analysis

Figure S6: Extended plot of figure 4b, showing the % lysogeny of polylysogens

Table S1. Linking genome IDs in figure 1a to IMG genome IDs – these can be found within the Integrated Microbial Genomes & Microbiomes (IMG/M) system at the DOE Joint Genome Institute (https://img.jgi.doe.gov/)

Table S2. Pairwise Wilcoxan rank-sum test and effect size for virulence of Phi3TΔ*aimP* against *B. subtilis* 168Δ6 in the presence of signals compared to no signal control, figure 2a. Significant results are highlight bold.

Table S3. Pairwise Wilcoxan rank-sum test results testing the difference in fluorescence of the reporter strain B. subtilis 168Δ6::aimR, p_aimX_-GFP (normalised to no signal control) in the presence of a panel of signal, compared to the no signal control, figure 1b. Significant results are highlight bold.

Table S4. Mutations in the signal peptides and how they could affect binding to Phi3T *aimR* to explain the differences in response of Phi3T lysis/lysogeny in the presence of these related signals.

Table S5. Pairwise Wilcoxan rank-sum test results for the efficiency of lysogeny of Phi3T in response to signals (figure 1c). Significant results are highlight bold.

Table S6. Pairwise Wilcoxan rank-sum results for prophage excision of Phi3T lysogen in response to signals, figure 1d. Significant results are highlight bold.

Table S7. Phi3T response to signals mixed linear model results

Table S8. Pairwise Wilcoxan rank-sum analysis of virulence of WT Phi3T compared to Phi3T.SIIRGA on host B. subtilis 168Δ6 in conditioned media (figure 4a). Significant results are highlighted in bold.

Table S9. Strains of bacteria used in this study

Table S10. Peptide table, clade and origin

Table S11. Primers used for modification of fluorescent plasmid Table S12. Mutant phages used in this study

Table S13. Primers, gblocks and conditions used to construct mutant phages Table S14. Strains, species and origin used in the host range assay

Table S15. Droplet digital PCR primers and PCR conditions

