## Supplementary data for "Arbitrium phages can manipulate each other’s lysis - lysogeny decisions"

Figure S1. The frequency of *Bacillus* genomes containing between 1 and 8 arbitrium-carrying prophages.

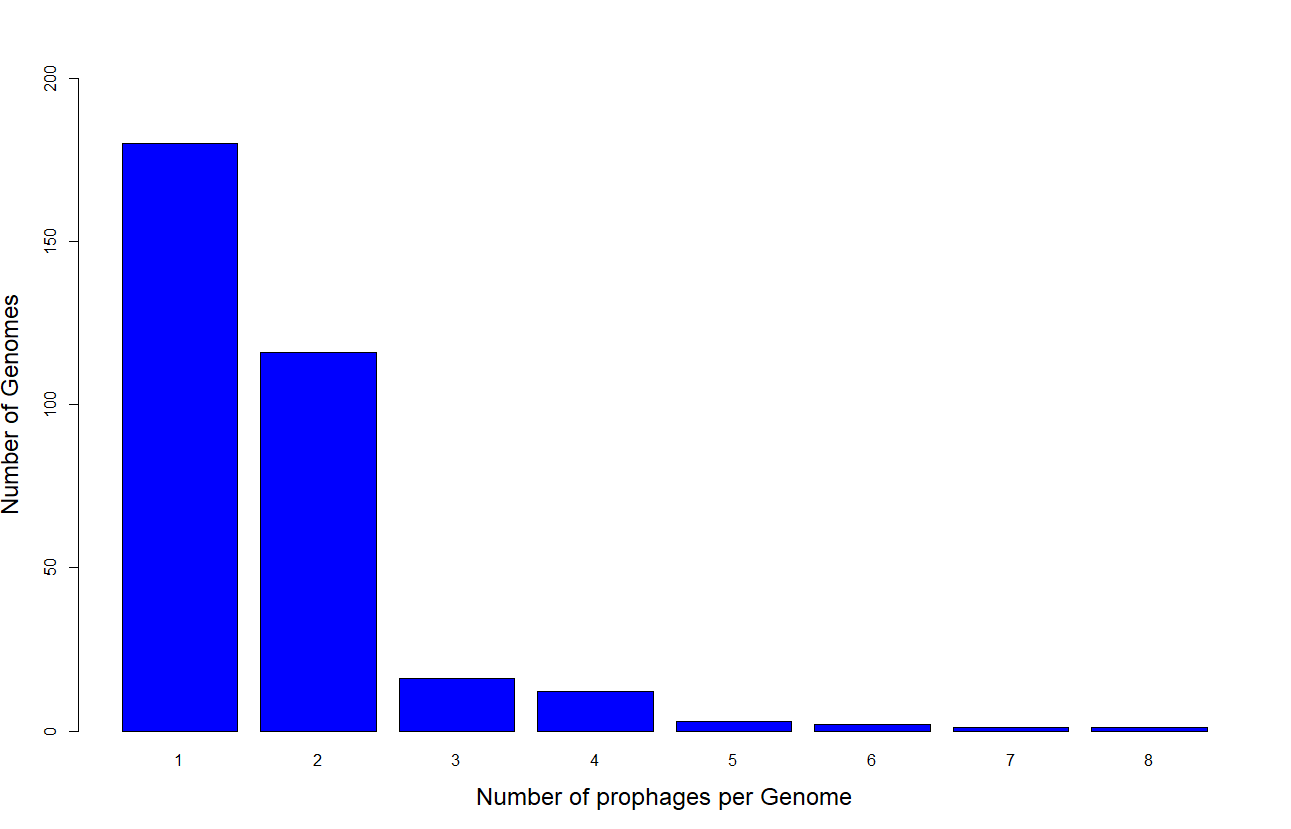

Figure S2. Virulence data calculated from growth curve OD_600_ data 16 SpBeta-like phages against a panel of 30 *Bacillus* hosts, used to calculate host range similarity (figure 1b).

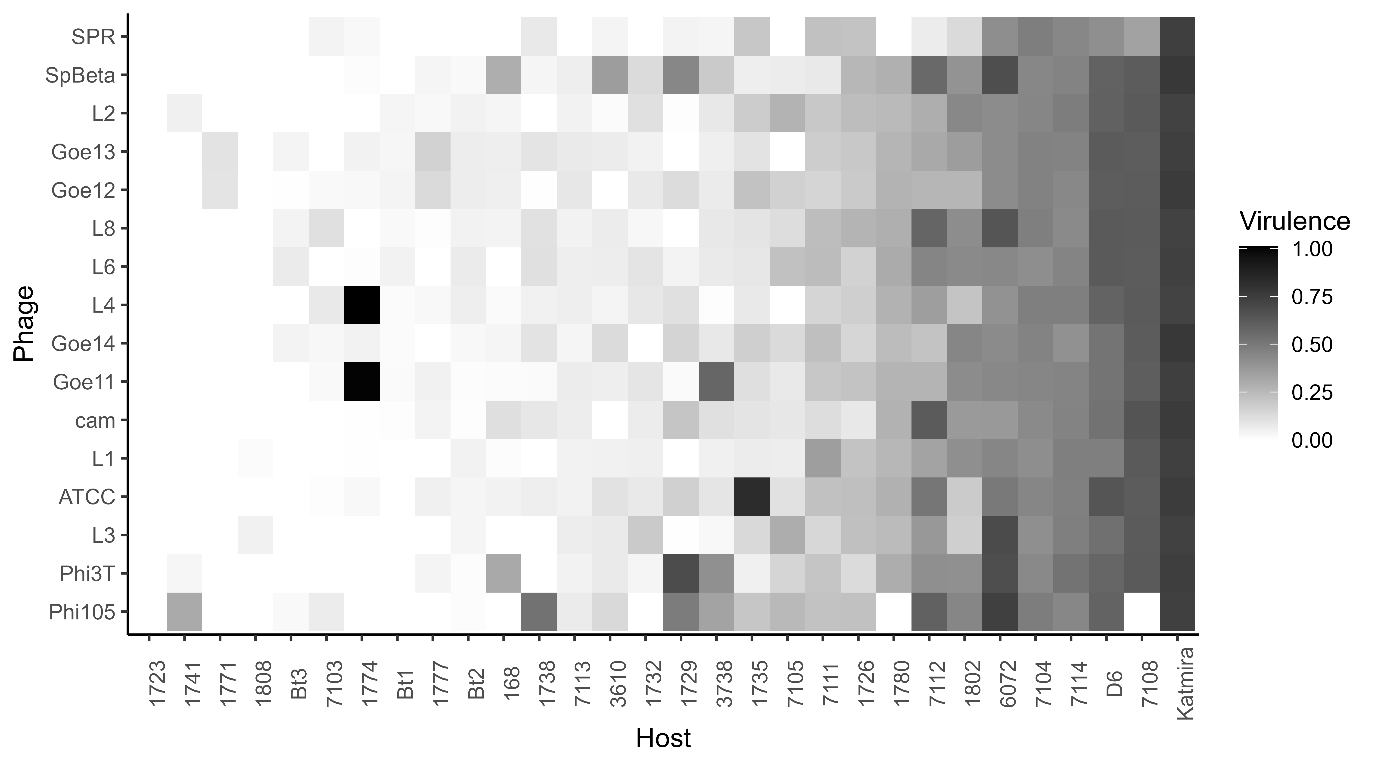

Figure S3. Intergenomic similarity (using Viridic (Moraru et al., 2020, <https://rhea.icbm.uni-oldenburg.de/viridic/>) between a collection of 15 phages carrying clade 2 arbitrium systems, the peptide signal each phage produces are grouped by colour.

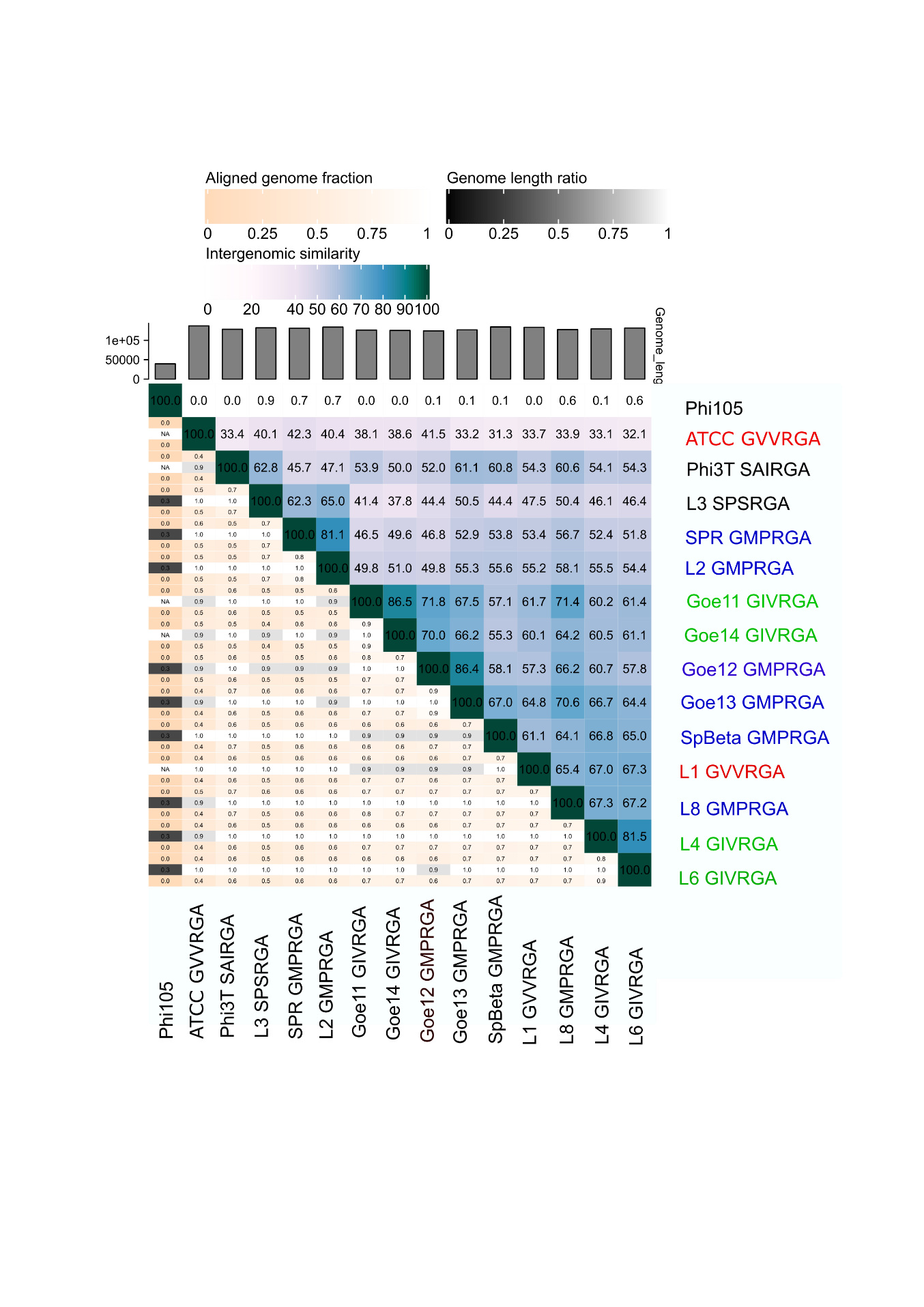

Figure S4: Maximum-likelihood phylogenetic trees (inferred using IQtree) for *aimR* and the large terminase (*terL*) gene from 398 arbitrium-carrying phage. The aimR tree contains reference samples to enable clustering by aimR clade.

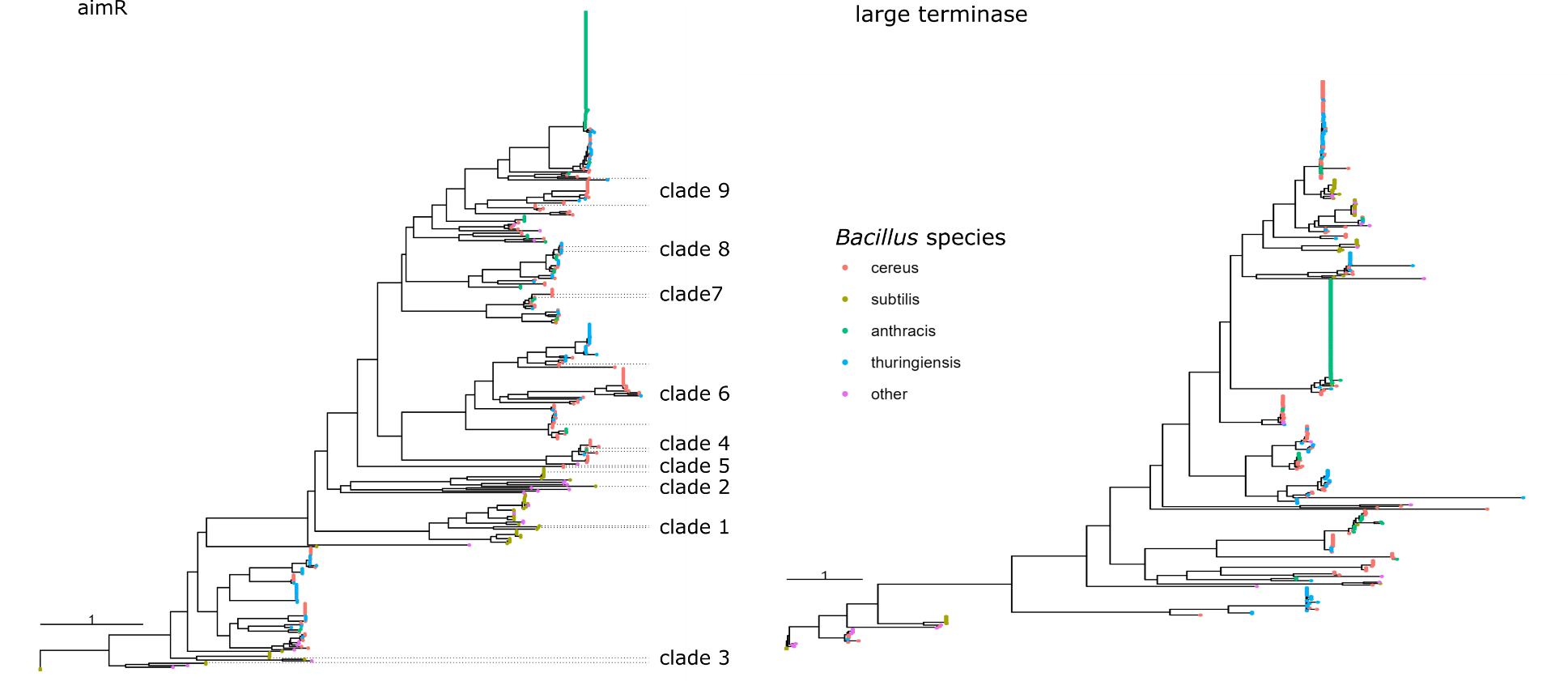

Figure S5. The virulence of WT Phi3T and Phi3T.SIIRGA phages against *B. subtilis 168Δ6* in media conditioned by eight phages that produce no signal (Phi3T ΔaimP and Phi3T ΔaimRPX), SAIRGA (WT Phi3T), SIIRGA (Phi3T.SIIRGA), GIVRGA (Goe11 and Goe14) and GMPRGA (Goe12 and Goe13).

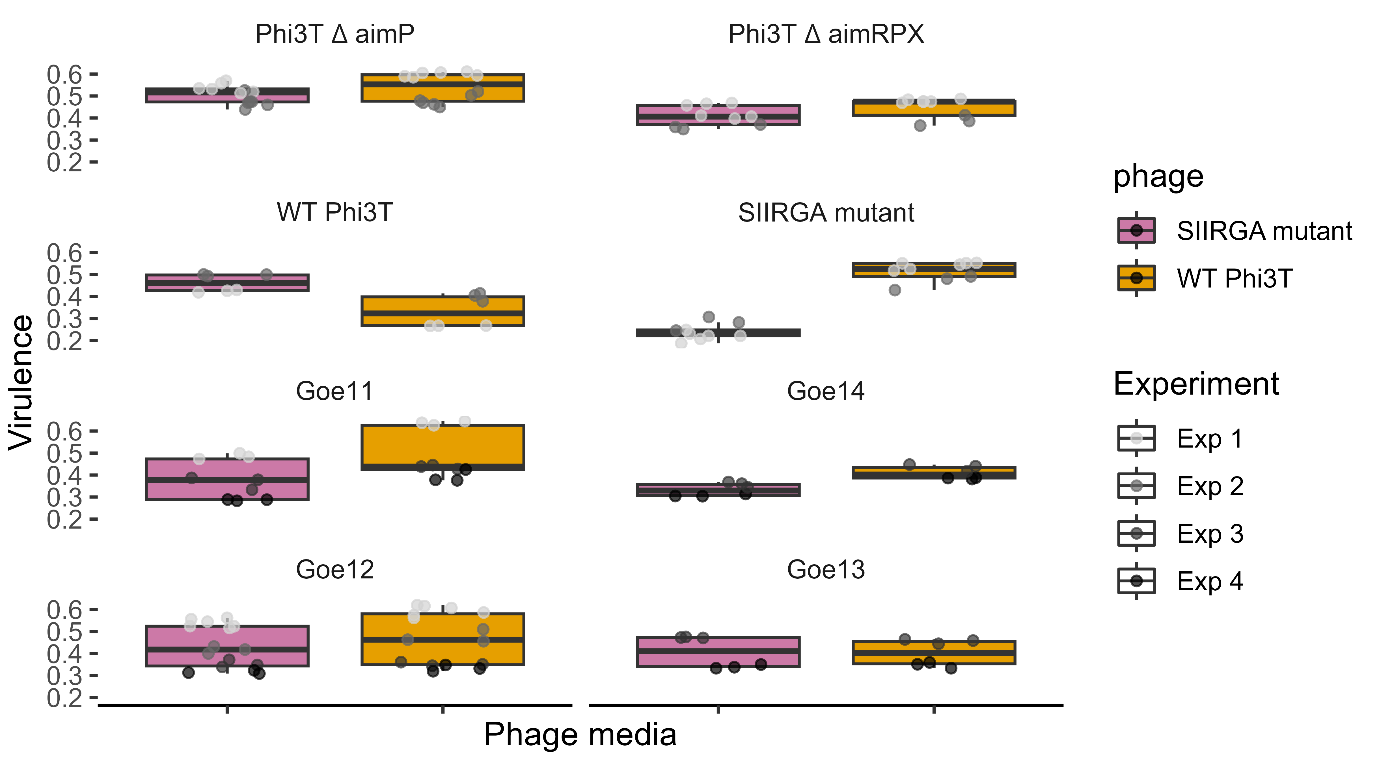

Figure S6. Percentage polylysogeny of *B. subtilis168Δ6* of WT Phi3T and Phi3T.SIIRGA during free phage co-infection with Goe11, at 2 hours post infection, showing that polylysogeny was rare.

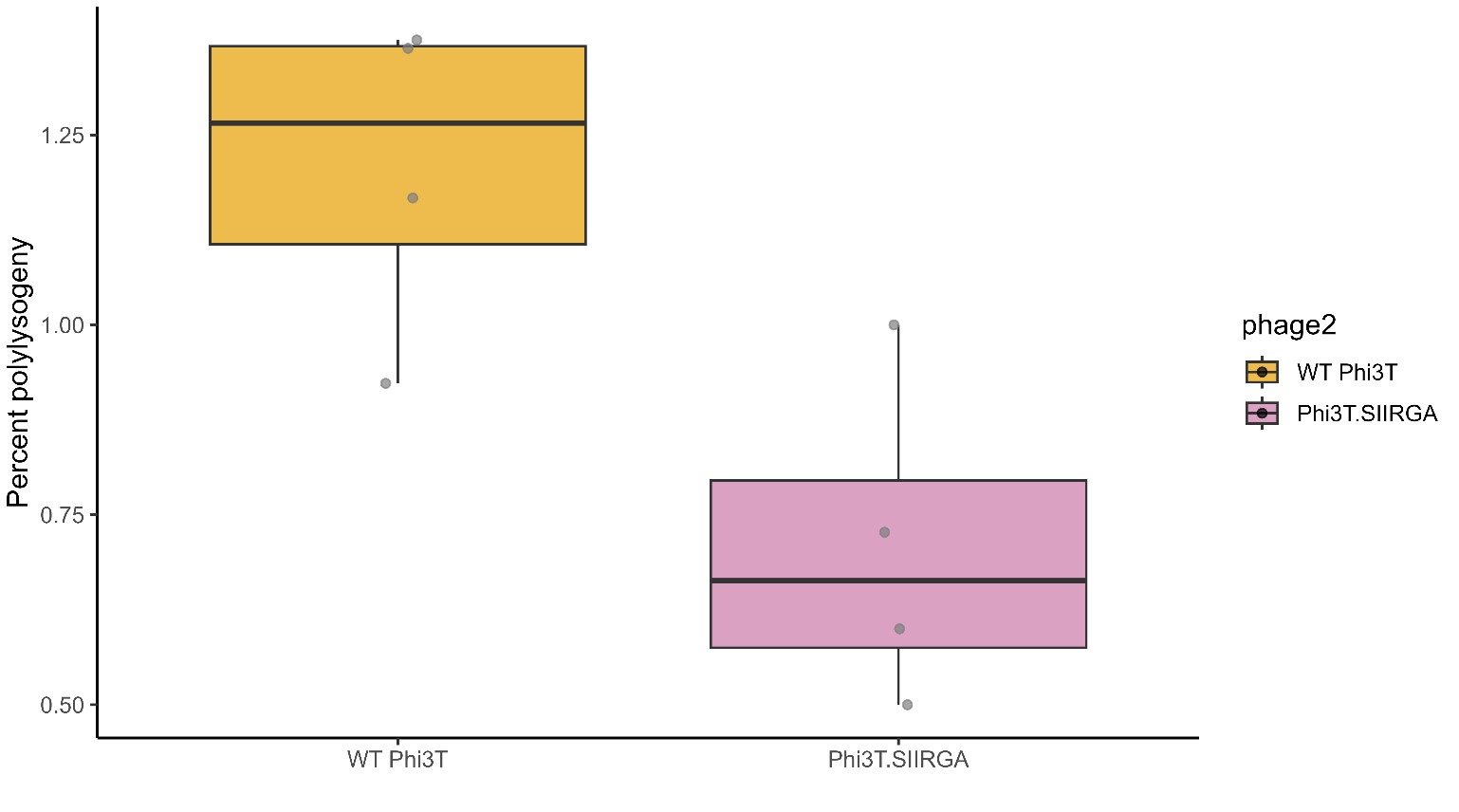

Table S1. The Integrated Microbial Genomes & Microbiomes (IMG/M) (<https://img.jgi.doe.gov/>) IDs for each genome, species, aimR clade and the ID used on in figure 1a.

| **IMG_genome_ID** | **species** | **aimR clade** | **Fig.1a genome ID** |
| --- | --- | --- | --- |
| 2634166487 | Bacillus_anthracis | 3 | Ba01 |
| 2634166487 | Bacillus_anthracis | 3 | Ba01 |
| 2634166487 | Bacillus_anthracis | 4 | Ba01 |
| 2634166487 | Bacillus_anthracis | 9 | Ba01 |
| 2597490031 | Bacillus_anthracis | 4 | Ba02 |
| 2597490031 | Bacillus_anthracis | 9 | Ba02 |
| 2597490031 | Bacillus_anthracis | 9 | Ba02 |
| unknown | Bacillus_anthracis | 4 | Ba03 |
| unknown | Bacillus_anthracis | 9 | Ba03 |
| unknown | Bacillus_anthracis | 9 | Ba03 |
| 2513237215 | Bacillus_anthracis | 4 | Ba04 |
| 2513237215 | Bacillus_anthracis | 9 | Ba04 |
| 2558860177 | Bacillus_anthracis | 4 | Ba05 |
| 2558860177 | Bacillus_anthracis | 9 | Ba05 |
| 2558860236 | Bacillus_anthracis | 4 | Ba06 |
| 2558860236 | Bacillus_anthracis | 9 | Ba06 |
| 2563366594 | Bacillus_anthracis | 4 | Ba07 |
| 2563366594 | Bacillus_anthracis | 9 | Ba07 |
| 2588253743 | Bacillus_anthracis | 4 | Ba08 |
| 2588253743 | Bacillus_anthracis | 9 | Ba08 |
| 2597489971 | Bacillus_anthracis | 4 | Ba09 |
| 2597489971 | Bacillus_anthracis | 9 | Ba09 |
| 2597489974 | Bacillus_anthracis | 4 | Ba10 |
| 2597489974 | Bacillus_anthracis | 9 | Ba10 |
| 2597489977 | Bacillus_anthracis | 4 | Ba11 |
| 2597489977 | Bacillus_anthracis | 9 | Ba11 |
| 2597490107 | Bacillus_anthracis | 4 | Ba12 |
| 2597490107 | Bacillus_anthracis | 9 | Ba12 |
| 2597490111 | Bacillus_anthracis | 4 | Ba13 |
| 2597490111 | Bacillus_anthracis | 9 | Ba13 |
| 2597490114 | Bacillus_anthracis | 4 | Ba14 |
| 2597490114 | Bacillus_anthracis | 9 | Ba14 |
| 2597490165 | Bacillus_anthracis | 4 | Ba15 |
| 2597490165 | Bacillus_anthracis | 9 | Ba15 |
| 2597490166 | Bacillus_anthracis | 4 | Ba16 |
| 2597490166 | Bacillus_anthracis | 9 | Ba16 |
| 2609460106 | Bacillus_anthracis | 4 | Ba17 |
| 2609460106 | Bacillus_anthracis | 9 | Ba17 |
| 2609460179 | Bacillus_anthracis | 4 | Ba18 |
| 2609460179 | Bacillus_anthracis | 9 | Ba18 |
| 2617271012 | Bacillus_anthracis | 4 | Ba19 |
| 2617271012 | Bacillus_anthracis | 9 | Ba19 |
| 2627853614 | Bacillus_anthracis | 4 | Ba20 |
| 2627853614 | Bacillus_anthracis | 9 | Ba20 |
| 2627853679 | Bacillus_anthracis | 4 | Ba21 |
| 2627853679 | Bacillus_anthracis | 9 | Ba21 |
| 2627853693 | Bacillus_anthracis | 4 | Ba22 |
| 2627853693 | Bacillus_anthracis | 9 | Ba22 |
| 2627853729 | Bacillus_anthracis | 4 | Ba23 |
| 2627853729 | Bacillus_anthracis | 9 | Ba23 |
| 2627853831 | Bacillus_anthracis | 4 | Ba24 |
| 2627853831 | Bacillus_anthracis | 9 | Ba24 |
| 2627854013 | Bacillus_anthracis | 4 | Ba25 |
| 2627854013 | Bacillus_anthracis | 9 | Ba25 |
| 2627854208 | Bacillus_anthracis | 4 | Ba26 |
| 2627854208 | Bacillus_anthracis | 9 | Ba26 |
| 2630968468 | Bacillus_anthracis | 4 | Ba27 |
| 2630968468 | Bacillus_anthracis | 9 | Ba27 |
| 2630968618 | Bacillus_anthracis | 4 | Ba28 |
| 2630968618 | Bacillus_anthracis | 9 | Ba28 |
| 2630968928 | Bacillus_anthracis | 4 | Ba29 |
| 2630968928 | Bacillus_anthracis | 9 | Ba29 |
| 2634166493 | Bacillus_anthracis | 4 | Ba30 |
| 2634166493 | Bacillus_anthracis | 9 | Ba30 |
| 2636415687 | Bacillus_anthracis | 4 | Ba31 |
| 2636415687 | Bacillus_anthracis | 9 | Ba31 |
| 2636415735 | Bacillus_anthracis | 4 | Ba32 |
| 2636415735 | Bacillus_anthracis | 9 | Ba32 |
| 2639762758 | Bacillus_anthracis | 4 | Ba33 |
| 2639762758 | Bacillus_anthracis | 9 | Ba33 |
| 2639762891 | Bacillus_anthracis | 4 | Ba34 |
| 2639762891 | Bacillus_anthracis | 9 | Ba34 |
| 2645727705 | Bacillus_anthracis | 4 | Ba35 |
| 2645727705 | Bacillus_anthracis | 9 | Ba35 |
| 2645728089 | Bacillus_anthracis | 4 | Ba36 |
| 2645728089 | Bacillus_anthracis | 9 | Ba36 |
| 2651869867 | Bacillus_anthracis | 4 | Ba37 |
| 2651869867 | Bacillus_anthracis | 9 | Ba37 |
| 2654588018 | Bacillus_anthracis | 4 | Ba38 |
| 2654588018 | Bacillus_anthracis | 9 | Ba38 |
| 2660238424 | Bacillus_anthracis | 4 | Ba39 |
| 2660238424 | Bacillus_anthracis | 9 | Ba39 |
| 2660238517 | Bacillus_anthracis | 4 | Ba40 |
| 2660238517 | Bacillus_anthracis | 9 | Ba40 |
| 2663763157 | Bacillus_anthracis | 4 | Ba41 |
| 2663763157 | Bacillus_anthracis | 9 | Ba41 |
| 2667527537 | Bacillus_anthracis | 4 | Ba42 |
| 2667527537 | Bacillus_anthracis | 9 | Ba42 |
| 2667527762 | Bacillus_anthracis | 4 | Ba43 |
| 2667527762 | Bacillus_anthracis | 9 | Ba43 |
| 2667527873 | Bacillus_anthracis | 4 | Ba44 |
| 2667527873 | Bacillus_anthracis | 9 | Ba44 |
| 2667527990 | Bacillus_anthracis | 4 | Ba45 |
| 2667527990 | Bacillus_anthracis | 9 | Ba45 |
| 2667528097 | Bacillus_anthracis | 4 | Ba46 |
| 2667528097 | Bacillus_anthracis | 9 | Ba46 |
| 2671180908 | Bacillus_anthracis | 4 | Ba47 |
| 2671180908 | Bacillus_anthracis | 9 | Ba47 |
| 2675903249 | Bacillus_anthracis | 4 | Ba48 |
| 2675903249 | Bacillus_anthracis | 9 | Ba48 |
| 2675903400 | Bacillus_anthracis | 4 | Ba49 |
| 2675903400 | Bacillus_anthracis | 9 | Ba49 |
| 2681813376 | Bacillus_anthracis | 4 | Ba50 |
| 2681813376 | Bacillus_anthracis | 9 | Ba50 |
| 2681813377 | Bacillus_anthracis | 4 | Ba51 |
| 2681813377 | Bacillus_anthracis | 9 | Ba51 |
| 2681813378 | Bacillus_anthracis | 4 | Ba52 |
| 2681813378 | Bacillus_anthracis | 9 | Ba52 |
| 2684622605 | Bacillus_anthracis | 4 | Ba53 |
| 2684622605 | Bacillus_anthracis | 9 | Ba53 |
| 2695420813 | Bacillus_anthracis | 4 | Ba54 |
| 2695420813 | Bacillus_anthracis | 9 | Ba54 |
| 2695420879 | Bacillus_anthracis | 4 | Ba55 |
| 2695420879 | Bacillus_anthracis | 9 | Ba55 |
| 2700989522 | Bacillus_anthracis | 4 | Ba56 |
| 2700989522 | Bacillus_anthracis | 9 | Ba56 |
| 2711768120 | Bacillus_anthracis | 4 | Ba57 |
| 2711768120 | Bacillus_anthracis | 9 | Ba57 |
| 2713897124 | Bacillus_anthracis | 4 | Ba58 |
| 2713897124 | Bacillus_anthracis | 9 | Ba58 |
| 2713897125 | Bacillus_anthracis | 4 | Ba59 |
| 2713897125 | Bacillus_anthracis | 9 | Ba59 |
| 2713897126 | Bacillus_anthracis | 4 | Ba60 |
| 2713897126 | Bacillus_anthracis | 9 | Ba60 |
| 2713897127 | Bacillus_anthracis | 4 | Ba61 |
| 2713897127 | Bacillus_anthracis | 9 | Ba61 |
| 2713897128 | Bacillus_anthracis | 4 | Ba62 |
| 2713897128 | Bacillus_anthracis | 9 | Ba62 |
| 2713897129 | Bacillus_anthracis | 4 | Ba63 |
| 2713897129 | Bacillus_anthracis | 9 | Ba63 |
| 2713897130 | Bacillus_anthracis | 4 | Ba64 |
| 2713897130 | Bacillus_anthracis | 9 | Ba64 |
| 2713897133 | Bacillus_anthracis | 4 | Ba65 |
| 2713897133 | Bacillus_anthracis | 9 | Ba65 |
| 2713897137 | Bacillus_anthracis | 4 | Ba66 |
| 2713897137 | Bacillus_anthracis | 9 | Ba66 |
| 2713897138 | Bacillus_anthracis | 4 | Ba67 |
| 2713897138 | Bacillus_anthracis | 9 | Ba67 |
| 637000013 | Bacillus_anthracis | 4 | Ba68 |
| 637000013 | Bacillus_anthracis | 9 | Ba68 |
| 637000015 | Bacillus_anthracis | 4 | Ba69 |
| 637000015 | Bacillus_anthracis | 9 | Ba69 |
| 641736155 | Bacillus_anthracis | 4 | Ba70 |
| 641736155 | Bacillus_anthracis | 9 | Ba70 |
| 641736173 | Bacillus_anthracis | 4 | Ba71 |
| 641736173 | Bacillus_anthracis | 9 | Ba71 |
| 643692005 | Bacillus_anthracis | 4 | Ba72 |
| 643692005 | Bacillus_anthracis | 9 | Ba72 |
| 643692006 | Bacillus_anthracis | 4 | Ba73 |
| 643692006 | Bacillus_anthracis | 9 | Ba73 |
| 2519103165 | Bacillus_cereus | 3 | Bc01 |
| 2519103165 | Bacillus_cereus | 6 | Bc01 |
| 2519103165 | Bacillus_cereus | 6 | Bc01 |
| 2519103165 | Bacillus_cereus | 9 | Bc01 |
| 2630968303 | Bacillus_cereus | 3 | Bc02 |
| 2630968303 | Bacillus_cereus | 4 | Bc02 |
| 2630968303 | Bacillus_cereus | 6 | Bc02 |
| 2511231081 | Bacillus_cereus | 3 | Bc03 |
| 2511231081 | Bacillus_cereus | 4 | Bc03 |
| 2511231081 | Bacillus_cereus | 7 | Bc03 |
| 2519103147 | Bacillus_cereus | 6 | Bc04 |
| 2519103147 | Bacillus_cereus | 6 | Bc04 |
| 2519103147 | Bacillus_cereus | 9 | Bc04 |
| 2519899844 | Bacillus_cereus | 7 | Bc05 |
| 2519899844 | Bacillus_cereus | 8 | Bc05 |
| 2519899844 | Bacillus_cereus | 9 | Bc05 |
| 2609460101 | Bacillus_cereus | 6 | Bc06 |
| 2609460101 | Bacillus_cereus | 6 | Bc06 |
| 2609460101 | Bacillus_cereus | 8 | Bc06 |
| 2519103125 | Bacillus_cereus | 6 | Bc07 |
| 2519103125 | Bacillus_cereus | 9 | Bc07 |
| 2519103149 | Bacillus_cereus | 7 | Bc08 |
| 2519103149 | Bacillus_cereus | 9 | Bc08 |
| 2531839360 | Bacillus_cereus | 3 | Bc09 |
| 2531839360 | Bacillus_cereus | 9 | Bc09 |
| 2531839574 | Bacillus_cereus | 6 | Bc10 |
| 2531839574 | Bacillus_cereus | 9 | Bc10 |
| 2534681696 | Bacillus_cereus | 8 | Bc11 |
| 2534681696 | Bacillus_cereus | 9 | Bc11 |
| 2537561734 | Bacillus_cereus | 4 | Bc12 |
| 2537561734 | Bacillus_cereus | 7 | Bc12 |
| 2537561846 | Bacillus_cereus | 6 | Bc13 |
| 2537561846 | Bacillus_cereus | 9 | Bc13 |
| 2537561852 | Bacillus_cereus | 3 | Bc14 |
| 2537561852 | Bacillus_cereus | 6 | Bc14 |
| 2537561853 | Bacillus_cereus | 9 | Bc15 |
| 2537561853 | Bacillus_cereus | 9 | Bc15 |
| 2537562207 | Bacillus_cereus | 3 | Bc16 |
| 2537562207 | Bacillus_cereus | 7 | Bc16 |
| 2537562214 | Bacillus_cereus | 7 | Bc17 |
| 2537562214 | Bacillus_cereus | 9 | Bc17 |
| 2597490033 | Bacillus_cereus | 3 | Bc18 |
| 2597490033 | Bacillus_cereus | 4 | Bc18 |
| 2627853723 | Bacillus_cereus | 8 | Bc19 |
| 2627853723 | Bacillus_cereus | 9 | Bc19 |
| 2627853898 | Bacillus_cereus | 8 | Bc20 |
| 2627853898 | Bacillus_cereus | 9 | Bc20 |
| 2636415615 | Bacillus_cereus | 8 | Bc21 |
| 2636415615 | Bacillus_cereus | 9 | Bc21 |
| 2684622616 | Bacillus_cereus | 6 | Bc22 |
| 2684622616 | Bacillus_cereus | 8 | Bc22 |
| 2703718890 | Bacillus_cereus | 7 | Bc23 |
| 2703718890 | Bacillus_cereus | 7 | Bc23 |
| 2728369135 | Bacillus_cereus | 3 | Bc24 |
| 2728369135 | Bacillus_cereus | 4 | Bc24 |
| 2728369139 | Bacillus_cereus | 4 | Bc25 |
| 2728369139 | Bacillus_cereus | 6 | Bc25 |
| 2728369141 | Bacillus_cereus | 7 | Bc26 |
| 2728369141 | Bacillus_cereus | 9 | Bc26 |
| 2728369164 | Bacillus_cereus | 6 | Bc27 |
| 2728369164 | Bacillus_cereus | 6 | Bc27 |
| 643348510 | Bacillus_cereus | 3 | Bc28 |
| 643348510 | Bacillus_cereus | 4 | Bc28 |
| 2561511152 | Bacillus_thuringiensis | 3 | Bt01 |
| 2561511152 | Bacillus_thuringiensis | 3 | Bt01 |
| 2561511152 | Bacillus_thuringiensis | 3 | Bt01 |
| 2561511152 | Bacillus_thuringiensis | 4 | Bt01 |
| 2561511152 | Bacillus_thuringiensis | 6 | Bt01 |
| 2561511152 | Bacillus_thuringiensis | 6 | Bt01 |
| 2561511152 | Bacillus_thuringiensis | 9 | Bt01 |
| 2561511152 | Bacillus_thuringiensis | 9 | Bt01 |
| 2645727773 | Bacillus_thuringiensis | 3 | Bt02 |
| 2645727773 | Bacillus_thuringiensis | 3 | Bt02 |
| 2645727773 | Bacillus_thuringiensis | 3 | Bt02 |
| 2645727773 | Bacillus_thuringiensis | 4 | Bt02 |
| 2645727773 | Bacillus_thuringiensis | 6 | Bt02 |
| 2645727773 | Bacillus_thuringiensis | 6 | Bt02 |
| 2645727773 | Bacillus_thuringiensis | 9 | Bt02 |
| 2521172680 | Bacillus_thuringiensis | 3 | Bt03 |
| 2521172680 | Bacillus_thuringiensis | 4 | Bt03 |
| 2521172680 | Bacillus_thuringiensis | 6 | Bt03 |
| 2521172680 | Bacillus_thuringiensis | 6 | Bt03 |
| 2521172680 | Bacillus_thuringiensis | 9 | Bt03 |
| 2521172680 | Bacillus_thuringiensis | 9 | Bt03 |
| 2687453486 | Bacillus_thuringiensis | 3 | Bt04 |
| 2687453486 | Bacillus_thuringiensis | 3 | Bt04 |
| 2687453486 | Bacillus_thuringiensis | 6 | Bt04 |
| 2687453486 | Bacillus_thuringiensis | 6 | Bt04 |
| 2687453486 | Bacillus_thuringiensis | 9 | Bt04 |
| 2687453486 | Bacillus_thuringiensis | 9 | Bt04 |
| 2671180266 | Bacillus_thuringiensis | 3 | Bt05 |
| 2671180266 | Bacillus_thuringiensis | 3 | Bt05 |
| 2671180266 | Bacillus_thuringiensis | 6 | Bt05 |
| 2671180266 | Bacillus_thuringiensis | 6 | Bt05 |
| 2671180266 | Bacillus_thuringiensis | 9 | Bt05 |
| 2654588007 | Bacillus_thuringiensis | 3 | Bt06 |
| 2654588007 | Bacillus_thuringiensis | 3 | Bt06 |
| 2654588007 | Bacillus_thuringiensis | 6 | Bt06 |
| 2654588007 | Bacillus_thuringiensis | 6 | Bt06 |
| 2654588007 | Bacillus_thuringiensis | 9 | Bt06 |
| 2597490277 | Bacillus_thuringiensis | 3 | Bt07 |
| 2597490277 | Bacillus_thuringiensis | 3 | Bt07 |
| 2597490277 | Bacillus_thuringiensis | 6 | Bt07 |
| 2609460077 | Bacillus_thuringiensis | 6 | Bt07 |
| 2609460077 | Bacillus_thuringiensis | 6 | Bt07 |
| 2636415423 | Bacillus_thuringiensis | 3 | Bt08 |
| 2636415423 | Bacillus_thuringiensis | 3 | Bt08 |
| 2636415423 | Bacillus_thuringiensis | 6 | Bt08 |
| 2636415423 | Bacillus_thuringiensis | 6 | Bt08 |
| 2636415423 | Bacillus_thuringiensis | 9 | Bt08 |
| 651053003 | Bacillus_thuringiensis | 3 | Bt09 |
| 651053003 | Bacillus_thuringiensis | 4 | Bt09 |
| 651053003 | Bacillus_thuringiensis | 6 | Bt09 |
| 651053003 | Bacillus_thuringiensis | 8 | Bt09 |
| 2585427662 | Bacillus_thuringiensis | 3 | Bt10 |
| 2585427662 | Bacillus_thuringiensis | 6 | Bt10 |
| 2585427662 | Bacillus_thuringiensis | 6 | Bt10 |
| 2585427662 | Bacillus_thuringiensis | 9 | Bt10 |
| 2561511195 | Bacillus_thuringiensis | 3 | Bt11 |
| 2561511195 | Bacillus_thuringiensis | 4 | Bt11 |
| 2561511195 | Bacillus_thuringiensis | 6 | Bt11 |
| 2561511195 | Bacillus_thuringiensis | 8 | Bt11 |
| 2561511198 | Bacillus_thuringiensis | 3 | Bt12 |
| 2561511198 | Bacillus_thuringiensis | 3 | Bt12 |
| 2561511198 | Bacillus_thuringiensis | 6 | Bt12 |
| 2561511198 | Bacillus_thuringiensis | 6 | Bt12 |
| 2630968894 | Bacillus_thuringiensis | 3 | Bt13 |
| 2630968894 | Bacillus_thuringiensis | 6 | Bt13 |
| 2630968894 | Bacillus_thuringiensis | 6 | Bt13 |
| 2630968894 | Bacillus_thuringiensis | 9 | Bt13 |
| 2687453290 | Bacillus_thuringiensis | 4 | Bt14 |
| 2687453290 | Bacillus_thuringiensis | 6 | Bt14 |
| 2687453290 | Bacillus_thuringiensis | 8 | Bt14 |
| 2687453290 | Bacillus_thuringiensis | 9 | Bt14 |
| 2521172701 | Bacillus_thuringiensis | 6 | Bt19 |
| 2521172701 | Bacillus_thuringiensis | 8 | Bt19 |
| 2521172701 | Bacillus_thuringiensis | 9 | Bt19 |
| 2521172701 | Bacillus_thuringiensis | 9 | Bt19 |
| 2522572172 | Bacillus_thuringiensis | 3 | Bt20 |
| 2522572172 | Bacillus_thuringiensis | 4 | Bt20 |
| 2522572172 | Bacillus_thuringiensis | 6 | Bt20 |
| 2522572172 | Bacillus_thuringiensis | 8 | Bt20 |
| 2654588002 | Bacillus_thuringiensis | 6 | Bt21 |
| 2654588002 | Bacillus_thuringiensis | 8 | Bt21 |
| 2654588002 | Bacillus_thuringiensis | 9 | Bt21 |
| 2630968597 | Bacillus_thuringiensis | 3 | Bt22 |
| 2630968597 | Bacillus_thuringiensis | 3 | Bt22 |
| 2630968597 | Bacillus_thuringiensis | 9 | Bt22 |
| 2518645577 | Bacillus_thuringiensis | 3 | Bt23 |
| 2518645577 | Bacillus_thuringiensis | 6 | Bt23 |
| 2518645577 | Bacillus_thuringiensis | 6 | Bt23 |
| 2627854229 | Bacillus_thuringiensis | 3 | Bt24 |
| 2627854229 | Bacillus_thuringiensis | 6 | Bt24 |
| 2627854229 | Bacillus_thuringiensis | 6 | Bt24 |
| 2627854070 | Bacillus_thuringiensis | 6 | Bt25 |
| 2627854070 | Bacillus_thuringiensis | 6 | Bt25 |
| 2627854070 | Bacillus_thuringiensis | 6 | Bt25 |
| 2545824726 | Bacillus_thuringiensis | 3 | Bt26 |
| 2545824726 | Bacillus_thuringiensis | 6 | Bt26 |
| 2576861478 | Bacillus_thuringiensis | 3 | Bt27 |
| 2576861478 | Bacillus_thuringiensis | 6 | Bt27 |
| 2654588052 | Bacillus_thuringiensis | 6 | Bt28 |
| 2654588052 | Bacillus_thuringiensis | 9 | Bt28 |
| 2684623201 | Bacillus_thuringiensis | 3 | Bt29 |
| 2684623201 | Bacillus_thuringiensis | 6 | Bt29 |
| 643886088 | Bacillus_thuringiensis | 7 | Bt30 |
| 643886088 | Bacillus_thuringiensis | 9 | Bt30 |
| 643886126 | Bacillus_thuringiensis | 6 | Bt31 |
| 643886126 | Bacillus_thuringiensis | 9 | Bt31 |
| 650377908 | Bacillus_thuringiensis | 6 | Bt32 |
| 650377908 | Bacillus_thuringiensis | 9 | Bt32 |
| 2540341129 | Bacillus_amyloliquefaciens | 1 | Bam01 |
| 2540341129 | Bacillus_amyloliquefaciens | 2 | Bam01 |
| 2540341129 | Bacillus_amyloliquefaciens | 2 | Bam01 |
| 2540341129 | Bacillus_amyloliquefaciens | 2 | Bam01 |
| 2511231180 | Bacillus_amyloliquefaciens | 2 | Bam02 |
| 2511231180 | Bacillus_amyloliquefaciens | 2 | Bam02 |
| 2511231180 | Bacillus_amyloliquefaciens | 3 | Bam02 |
| 651053002 | Bacillus_amyloliquefaciens | 2 | Bam03 |
| 651053002 | Bacillus_amyloliquefaciens | 2 | Bam03 |
| 651053002 | Bacillus_amyloliquefaciens | 3 | Bam03 |
| 2634166398 | Bacillus_amyloliquefaciens | 2 | Bam04 |
| 2634166398 | Bacillus_amyloliquefaciens | 3 | Bam04 |
| 2636415949 | Bacillus_amyloliquefaciens | 1 | Bam05 |
| 2636415949 | Bacillus_amyloliquefaciens | 3 | Bam05 |
| 2654587976 | Bacillus_amyloliquefaciens | 2 | Bam06 |
| 2654587976 | Bacillus_amyloliquefaciens | 2 | Bam06 |
| 649633008 | Bacillus_amyloliquefaciens | 2 | Bam07 |
| 649633008 | Bacillus_amyloliquefaciens | 3 | Bam07 |
| 651053001 | Bacillus_amyloliquefaciens | 2 | Bam08 |
| 651053001 | Bacillus_amyloliquefaciens | 2 | Bam08 |
| 642791616 | Bacillus_pumilus | 1 | Bp01 |
| 642791616 | Bacillus_pumilus | 3 | Bp01 |
| 2724679183 | Bacillus_nakamurai | 1 | Bn01 |
| 2724679183 | Bacillus_nakamurai | 2 | Bn01 |
| 2615840647 | Bacillus_methylotrophicus | 1 | Bm01 |
| 2615840647 | Bacillus_methylotrophicus | 2 | Bm01 |
| 2627853700 | Bacillus_subtilis | 2 | Bs01 |
| 2627853700 | Bacillus_subtilis | 2 | Bs02 |
| 2627853700 | Bacillus_subtilis | 3 | Bs03 |
| 2513237190 | Bacillus_velezensis | 1 | Bv01 |
| 2513237190 | Bacillus_velezensis | 2 | Bv01 |
| 2513237190 | Bacillus_velezensis | 2 | Bv01 |
| 2513237190 | Bacillus_velezensis | 2 | Bv01 |
| 2511231119 | Bacillus_velezensis | 0 | Bv02 |
| 2511231119 | Bacillus_velezensis | 1 | Bv02 |
| 2576861459 | Bacillus_velezensis | 1 | Bv03 |
| 2576861459 | Bacillus_velezensis | 2 | Bv03 |
| 2627853654 | Bacillus_sp | 7 | Bsp01 |
| 2627853654 | Bacillus_sp | 8 | Bsp01 |
| 2627853968 | Bacillus_sp | 7 | Bsp02 |
| 2627853968 | Bacillus_sp | 8 | Bsp02 |
| 2643221729 | Bacillus_sp | 6 | Bsp03 |
| 2643221729 | Bacillus_sp | 6 | Bsp03 |
| 2643221730 | Bacillus_sp | 3 | Bsp04 |
| 2643221730 | Bacillus_sp | 6 | Bsp04 |
| 2643221950 | Bacillus_sp | 2 | Bsp05 |
| 2643221950 | Bacillus_sp | 2 | Bsp05 |
| 2684623176 | Bacillus_sp | 2 | Bsp06 |
| 2684623176 | Bacillus_sp | 2 | Bsp06 |

Table S2. Pairwise Wilcoxan rank-sum test for virulence of Phi3T*ΔaimP* against *B. subtilis* 168Δ6 in the presence of each of 30 synthetic signal peptides compared to the no signal control (figure 2a). n1 = number of samples in no signal control group, n2 = number of samples in treatment group, W = test statistic, p = p-value, p.adjusted = p value adjusted for multiple comparisons (Bonferroni correction), Wilcoxan effect size = calculated using Coin package (>0.5 = strong effect). Signals yielding a significant result are highlighted bold.

| **group1** | **group2** | **n1** | **n2** | **W** | **p** | **p.adjusted** | **effect size** |
| --- | --- | --- | --- | --- | --- | --- | --- |
| No signal | AIGNGG | 9 | 9 | 47 | 0.605 | 1 | 0.135285 |
| No signal | AMGNGG | 9 | 9 | 60 | 0.094 | 1 | 0.405854 |
| No signal | DPPVGM | 9 | 9 | 53 | 0.297 | 1 | 0.260163 |
| No signal | DVRPGG | 9 | 9 | 46 | 0.666 | 1 | 0.114472 |
| No signal | EIKPGG | 9 | 9 | 50.5 | 0.401 | 1 | 0.208238 |
| No signal | EIRVGM | 9 | 9 | 53 | 0.297 | 1 | 0.260163 |
| No signal | GFGHGA | 9 | 9 | 58 | 0.136 | 1 | 0.364228 |
| No signal | GFGRGA | 9 | 9 | 35 | 0.666 | 1 | 0.114472 |
| No signal | GFPRGA | 9 | 9 | 52 | 0.34 | 1 | 0.23935 |
| No signal | GFTVGA | 9 | 9 | 54 | 0.258 | 1 | 0.280976 |
| No signal | GIVRGA | 9 | 3 | 27 | 0.009 | 0.27 | 0.720577 |
| No signal | GMPRGA | 9 | 9 | 59 | 0.113 | 1 | 0.385041 |
| No signal | GTGTRP | 9 | 9 | 57 | 0.161 | 1 | 0.343415 |
| **No signal** | **GVVRGA** | **9** | **6** | **54** | **4e-04** | **0.012** | **0.821584** |
| No signal | LMVDPGTH | 9 | 9 | 47 | 0.605 | 1 | 0.135285 |
| No signal | LRMDPGTGIG | 9 | 9 | 61 | 0.077 | 1 | 0.426667 |
| No signal | MEADPGGTGG | 9 | 9 | 54 | 0.258 | 1 | 0.280976 |
| No signal | MEVGPGGGMG | 9 | 9 | 38 | 0.863 | 1 | 0.052033 |
| No signal | MKMDPGTLG | 9 | 9 | 58 | 0.136 | 1 | 0.364228 |
| No signal | MMSEPGGGGW | 9 | 9 | 45 | 0.73 | 1 | 0.093659 |
| No signal | MMTDPGGS | 9 | 9 | 54 | 0.258 | 1 | 0.280976 |
| No signal | MMVDPGGGW | 9 | 9 | 59 | 0.113 | 1 | 0.385041 |
| No signal | MRMDPGTIG | 9 | 9 | 49 | 0.489 | 1 | 0.176911 |
| No signal | NPGRGA | 9 | 9 | 57 | 0.161 | 1 | 0.343415 |
| **No signal** | **SAIRGA** | **9** | **9** | **81** | **4.11e-05** | **0.001233** | **0.842927** |
| **No signal** | **SASRGA** | **9** | **9** | **81** | **4.11e-05** | **0.001233** | **0.842927** |
| **No signal** | **SIIRGA** | **9** | **9** | **81** | **4.11e-05** | **0.001233** | **0.842927** |
| **No signal** | **SPSRGA** | **9** | **9** | **81** | **4.11e-05** | **0.001233** | **0.842927** |
| No signal | TIGRGG | 9 | 9 | 55 | 0.222 | 1 | 0.301789 |
| No signal | YMIDPGPGGG | 9 | 9 | 50 | 0.436 | 1 | 0.197724 |

Table S3. Pairwise Wilcoxan rank-sum test results testing the difference in fluorescence of the reporter strain *B. subtilis 168Δ6::RX-GFP* (normalised to no signal control) in the presence of a panel of 30 signals, compared to the no signal control (figure 2b). Significant results are highlight bold.

| **group1** | **group2** | **n1** | **n2** | **W** | **p** | **p.adjusted** | **effect size** |
| --- | --- | --- | --- | --- | --- | --- | --- |
| No signal | AIGNGG | 9 | 6 | 28 | 0.949 | 1 | 0.033005 |
| No signal | AMGNGG | 9 | 6 | 20 | 0.435 | 1 | 0.216912 |
| No signal | DPPVGM | 9 | 6 | 35 | 0.319 | 1 | 0.274629 |
| No signal | DVRPGG | 9 | 6 | 36 | 0.277 | 1 | 0.297044 |
| No signal | EIKPGG | 9 | 6 | 28 | 0.949 | 1 | 0.033005 |
| No signal | EIRVGM | 9 | 6 | 28 | 0.949 | 1 | 0.033005 |
| No signal | GFGHGA | 9 | 6 | 28 | 0.949 | 1 | 0.033005 |
| No signal | GFGRGA | 9 | 6 | 30 | 0.756 | 1 | 0.096225 |
| No signal | GFPRGA | 9 | 6 | 28 | 0.949 | 1 | 0.033005 |
| No signal | GFTVGA | 9 | 6 | 39 | 0.126 | 1 | 0.411943 |
| No signal | GIVRGA | 9 | 6 | 32 | 0.565 | 1 | 0.165025 |
| No signal | GMPRGA | 9 | 9 | 43 | 0.846 | 1 | 0.057123 |
| No signal | GTGTRP | 9 | 6 | 32 | 0.565 | 1 | 0.165025 |
| No signal | **GVVRGA** | **9** | **6** | **0** | **0.002** | **0.051** | **0.829019** |
| No signal | LMVDPGTH | 9 | 6 | 26 | 0.95 | 1 | 0.032075 |
| No signal | LRMDPGTGIG | 9 | 6 | 36 | 0.277 | 1 | 0.297044 |
| No signal | MEADPGGTGG | 9 | 6 | 25 | 0.852 | 1 | 0.06415 |
| No signal | MEVGPGGGMG | 9 | 6 | 28 | 0.949 | 1 | 0.033005 |
| No signal | MKMDPGTLG | 9 | 6 | 42 | 0.042 | 1 | 0.543469 |
| No signal | MMSEPGGGGW | 9 | 6 | 35 | 0.319 | 1 | 0.274629 |
| No signal | MMTDPGGS | 9 | 6 | 23 | 0.674 | 1 | 0.12395 |
| No signal | MMVDPGGGW | 9 | 6 | 42 | 0.042 | 1 | 0.543469 |
| No signal | MRMDPGTIG | 9 | 6 | 39 | 0.126 | 1 | 0.411943 |
| No signal | NPGRGA | 9 | 6 | 35 | 0.319 | 1 | 0.274629 |
| No signal | **SAIRGA** | **9** | **9** | **0** | **0.000385** | **0.012** | **0.847311** |
| No signal | SASRGA | 9 | 6 | 0 | 0.002 | 0.051 | 0.829019 |
| No signal | **SIIRGA** | **9** | **9** | **0** | **0.000385** | **0.012** | **0.847311** |
| No signal | **SPSRGA** | **9** | **9** | **0** | **0.000385** | **0.012** | **0.847311** |
| No signal | TIGRGG | 9 | 6 | 42 | 0.042 | 1 | 0.543469 |
| No signal | YMIDPGPGGG | 9 | 6 | 42 | 0.042 | 1 | 0.543469 |

Table S4. Phi3T aimR responds to four non-cognate signals (SIIRGA, SASRGA, SPSRGA and GVVRGA) to varying degrees but not to GMPRGA, here we identify the amino acid substitutions in comparison to the cognate peptide (SAIRGA), and the effect each mutation has on the biochemistry of the peptide, which could affect their ability to bind to Phi3T AimR.

| **Peptide amino acid sequence** | **Mutation away from cognate** | **Effect of mutation** |
| --- | --- | --- |
| SAIRGA | Phi3T cognate signal |  |
| S**I**IRGA | Second pos: A to I | Mild, both hydrophobic |
| SA**S**RGA | Third pos: I to S | Changes sidechain from hydrophobic to polar |
| S**PS**RGA | Third pos: I to S  Second pos: A to P | As above.  The I to S appears to be more important than A to P, as both SASRGA and SPSRGA peptides produce a similar response |
| **GVV**RGA | First pos: S to G  Second pos: A to VThird pos: I to V | S to G removes the polar OH side chain associated with S.  Milder compared to A to I or A to M, as side chains become increasingly long from V to I to M.  Mild mutation. V has shorter sidechain but chemically similar to I. |
| **GMP**RGA | First pos: S to G  Second pos: A to M  Third pos: I to P | As above  Side chains of M are long compared to A  P has a ring structure and thus this is a more disruptive mutation compared to I to V |

Table S5. Pairwise Wilcoxan rank-sum test results for the efficiency of lysogeny of Phi3T of *B. subtilis 168Δ6* in response to synthetic signals (figure 2c). Significant results are highlight bold.

| **group1** | **group2** | **n1** | **n2** | **W** | **p** | **p.adjusted** | **effect size** |
| --- | --- | --- | --- | --- | --- | --- | --- |
| No signal | DVRPGG | 21 | 8 | 92.5 | 0.696 | 1 | 0.07718 |
| **No signal** | **GIVRGA** | **21** | **9** | **163** | **0.002** | **0.022** | **0.566937** |
| No signal | GMPRGA | 21 | 12 | 145 | 0.488 | 1 | 0.123976 |
| **No signal** | **GVVRGA** | **21** | **12** | **0** | **2.6e-06** | **2.86e-05** | **0.821469** |
| No signal | LMVDPGTH | 21 | 8 | 82.5 | 0.961 | 1 | 0.013608 |
| No signal | MMSEPGGGGW | 21 | 11 | 111.5 | 0.889 | 1 | 0.028109 |
| No signal | MMTDPGGS | 21 | 11 | 69.5 | 0.07 | 0.77 | 0.323198 |
| **No signal** | **SAIRGA** | **21** | **20** | **0** | **4.58e-08** | **5.038e-07** | **0.855846** |
| **No signal** | **SASRGA** | **21** | **12** | **0** | **2.6e-06** | **2.86e-05** | **0.821469** |
| **No signal** | **SIIRGA** | **21** | **20** | **49.5** | **2.97e-05** | **0.000327** | **0.65414** |
| **No signal** | **SPSRGA** | **21** | **12** | **0** | **2.61e-06** | **2.871e-05** | **0.821263** |

Table S6. Pairwise Wilcoxan rank-sum results for prophage excision of *B. subtilis 168Δ6*::Phi3T*Δaimp* lysogen in response to synthetic signals (figure 2d). Significant results are highlight bold

| **group1** | **group2** | **n1** | **n2** | **statistic** | **p** | **p.adjusted** | **Effect size** |
| --- | --- | --- | --- | --- | --- | --- | --- |
| **No signal** | **DVRPGG** | **12** | **12** | **144** | **3.56e-05** | **0.000392** | **0.849822** |
| No signal | GIVRGA | 12 | 12 | 101 | 0.095 | 1 | 0.347091 |
| No signal | GMPRGA | 12 | 12 | 112 | 0.021 | 0.231 | 0.476614 |
| **No signal** | **GVVRGA** | **12** | **12** | **141** | **7.41e-05** | **0.000815** | **0.814768** |
| **No signal** | **LMVDPGTH** | **12** | **12** | **126** | **0.002** | **0.022** | **0.639041** |
| **No signal** | **MMSEPGGGGW** | **12** | **12** | **143** | **4.52e-05** | **0.000497** | **0.838568** |
| No signal | MMTDPGGS | 12 | 12 | 93 | 0.23 | 1 | 0.25078 |
| **No signal** | **SAIRGA** | **12** | **12** | **144** | **3.48e-05** | **0.000383** | **0.850936** |
| **No signal** | **SASRGA** | **12** | **12** | **144** | **3.45e-05** | **0.00038** | **0.851309** |
| **No signal** | **SIIRGA** | **12** | **12** | **144** | **3.42e-05** | **0.000376** | **0.851681** |
| **No signal** | **SPSRGA** | **12** | **12** | **142** | **5.87e-05** | **0.000646** | **0.826036** |

Table S7. Ten natural (Goe11, Goe12, Goe13, Goe14, ATCC13952, Spbeta, L1 and L6) and mutant phages (Phi3T.SIIRGA and Phi3T.SASRGA) response to signals, linear mixed models analysis; a) full model b) Tukey pairwise comparison of each signal to no-signal.

a)

| **Phage** | **Chi-squared** | **deg.of freedom** | **P- value** |
| --- | --- | --- | --- |
| Goe12 | 247.91 | 7 | < 0.01 |
| Goe11 | 86.153 | 7 | < 0.01 |
| Goe13 | 76.495 | 7 | < 0.01 |
| ATTC13952 | 94.371 | 7 | < 0.01 |
| Phi3T.SIIRGA | 178.69 | 7 | < 0.01 |
| Phi3T | 124.47 | 7 | < 0.01 |
| SpBeta | 44.91 | 7 | < 0.01 |
| L1 | 148.98 | 7 | < 0.01 |
| L6 | 34.504 | 7 | < 0.01 |
| Phi3T.SASRGA | 24.218 | 7 | < 0.01 |

b)

| **Phage** | **Treatment** | **Tukey-adjusted P value (Comparison to No signal)** |
| --- | --- | --- |
| Goe12 | GIVRGA | <.0001 |
|  | GMPRGA | <.0001 |
|  | GVVRGA | 0.0574 |
|  | SAIRGA | 0.1414 |
|  | SASRGA | 0.255 |
|  | SIIRGA | 0.1314 |
|  | SPSRGA | 0.0063 |
| Goe11 | GIVRGA | 0.0001 |
|  | GMPRGA | 1 |
|  | GVVRGA | <.0001 |
|  | SAIRGA | 1 |
|  | SASRGA | 1 |
|  | SIIRGA | <.0001 |
|  | SPSRGA | 0.9988 |
| Goe13 | GIVRGA | <.0001 |
|  | GMPRGA | <.0001 |
|  | GVVRGA | 0.002 |
|  | SAIRGA | 0.9665 |
|  | SASRGA | 0.6356 |
|  | SIIRGA | 0.6059 |
|  | SPSRGA | 0.2355 |
| Phi3T | GIVRGA | 1 |
|  | GMPRGA | 1 |
|  | GVVRGA | <.0001 |
|  | SAIRGA | <.0001 |
|  | SASRGA | <.0001 |
|  | SIIRGA | <.0001 |
|  | SPSRGA | <.0001 |
| ATCC13952 | GIVRGA | <.0001 |
|  | GMPRGA | 0.9551 |
|  | GVVRGA | <.0001 |
|  | SAIRGA | 0.1486 |
|  | SASRGA | 1 |
|  | SIIRGA | 0.0002 |
|  | SPSRGA | 0.0701 |
| Phi3T.SIIRGA | GIVRGA | <.0001 |
|  | GMPRGA | 0.2145 |
|  | GVVRGA | <.0001 |
|  | SAIRGA | <.0001 |
|  | SASRGA | 0.9998 |
|  | SIIRGA | <.0001 |
|  | SPSRGA | 0.9738 |
| SpBeta | GIVRGA | 0.6751 |
|  | GMPRGA | <.0001 |
|  | GVVRGA | 0.119 |
|  | SAIRGA | 0.8104 |
|  | SASRGA | 0.7716 |
|  | SIIRGA | 1 |
|  | SPSRGA | 0.2872 |
| L1 | GIVRGA | <.0001 |
|  | GMPRGA | 0.9999 |
|  | GVVRGA | <.0001 |
|  | SAIRGA | 0.9843 |
|  | SASRGA | 0.9121 |
|  | SIIRGA | 0.0067 |
|  | SPSRGA | <.0001 |
| L6 | GIVRGA | 0.0002 |
|  | GMPRGA | 0.0469 |
|  | GVVRGA | 0.008 |
|  | SAIRGA | 0.9941 |
|  | SASRGA | 0.1604 |
|  | SIIRGA | 0.0023 |
|  | SPSRGA | 0.0129 |
| Phi3T.SASRGA | GIVRGA | 0.1796 |
|  | GMPRGA | 0.1313 |
|  | GVVRGA | 0.1868 |
|  | SAIRGA | 0.1867 |
|  | SASRGA | 0.0109 |
|  | SIIRGA | 0.8649 |
|  | SPSRGA | 0.0007 |

Table S8. Pairwise Wilcoxan rank-sum analysis of the virulence of WT Phi3T compared to Phi3T.SIIRGA on host *B. subtilis168Δ6* in eight conditioned medias (figure 4a). Significant results are highlighted in bold.

| **Phage_media** | **.y.** | **group1** | **group2** | **n1** | **n2** | **W** | **p** | **p.adjusted** |
| --- | --- | --- | --- | --- | --- | --- | --- | --- |
| Phi3TΔ*aimP* | virulence | Phi3T.SIIRGA | WT Phi3T | 12 | 12 | 50 | 0.219 | 1 |
| Phi3TΔ*aimRPX* | virulence | Phi3T.SIIRGA | WT Phi3T | 9 | 9 | 16 | 0.032 | 0.256 |
| **WT Phi3T** | **virulence** | **Phi3T.SIIRGA** | **WT Phi3T** | **6** | **6** | **36** | **0.002** | **0.016** |
| **Phi3T.SIIRGA** | **virulence** | **Phi3T.SIIRGA** | **WT Phi3T** | **9** | **9** | **0** | **4.11e-05** | **0.000329** |
| Goe11 | virulence | Phi3T.SIIRGA | WT Phi3T | 9 | 9 | 22 | 0.113 | 0.904 |
| **Goe14** | **virulence** | **Phi3T.SIIRGA** | **WT Phi3T** | **6** | **6** | **0** | **0.002** | **0.016** |
| Goe12 | virulence | Phi3T.SIIRGA | WT Phi3T | 15 | 15 | 85 | 0.267 | 1 |
| Goe13 | virulence | Phi3T.SIIRGA | WT Phi3T | 6 | 6 | 20 | 0.818 | 1 |

Table S9. Bacteria strains used in this study

| Strain | Origin | Details |
| --- | --- | --- |
| *B. subtilis* 168 Δ6 | Gifted from Jose Penades | Strain 168 with MGEs removed |
| *B. subtilis*168 Δ6 *amyE*::*aimR, p_aimX_-GFP* | Plasmid pDR111::aimRPX-GFP gifted by Gil Amitai (Sorek lab) | Modified in this study to remove *aimP* and transformed into *B. subtilis* 168Δ6: A reporter strain to determine the affinity of Phi3T aimR to non-cognate signal peptides |
| Bacillus subtilis BEST7003:*aimX* | Described previously, Erez et al, 2017. Gifted from the Sorek lab | *aimX* under a xylose inducible promoter to enable accurate PFU counts of lysogenic Phi3T |

| Arbitrium peptide | aimR clade | origin |
| --- | --- | --- |
| DPPVGM | 1 | Peptide 2.0 |
| EIRVGM | 1 | Peptide 2.0 |
| AMGNGG | 2 | Peptide 2.0 |
| GFTVGA | 2 | Peptide 2.0 |
| GFGHGA | 2 | Peptide 2.0 |
| AIGNGG | 2 | Peptide 2.0 |
| SAIRGA | 2 | ISCA Biochemicals Ltd |
| SIIRGA | 2 | Peptide 2.0 |
| SASRGA | 2 | Peptide 2.0 |
| SPSRGA | 2 | Peptide 2.0 |
| GMPRGA | 2 | Peptide 2.0 |
| NPGRGA | 2 | Peptide 2.0 |
| GTGTRP | 2 | Peptide 2.0 |
| TIGRGG | 2 | Peptide 2.0 |
| GFPRGA | 2 | Peptide 2.0 |
| GVVRGA | 2 | GenScript |
| GFGRGA | 2 | Peptide 2.0 |
| GIVRGA | 2 | Peptide 2.0 |
| EIKPGG | 3 | Peptide 2.0 |
| DVRPGG | 3 | Peptide 2.0 |
| MKMDPGTLG | 4 | Peptide 2.0 |
| MRMDPGTIG | 4 | Peptide 2.0 |
| MMTDPGGS | 6 | Peptide 2.0 |
| LMVDPGTH | 6 | Peptide 2.0 |
| MEVGPGGGMG | 7 | Peptide 2.0 |
| LRMDPGTGIG | 7 | Peptide 2.0 |
| MEADPGGTGG | 8 | Peptide 2.0 |
| YMIDPGPGGG | 8 | Peptide 2.0 |
| MMSEPGGGGW | 9 | Peptide 2.0 |
| MMVDPGGGW | 9 | Peptide 2.0 |

Table S10. Synthetic peptide sequences, the aimR clade they belong to and source company.

Table S11. Primers and PCR conditions used for modification of the fluorescent reporter plasmid pDR111_*aimRPX* - using the pDR111_*aimRPX* vector DNA as a template, fragments upstream (*aimR*) and downstream (*aimX*) of the aimP gene were amplified using the primers below

| **Target_primer** | **Primers** | **Amplification program** | **Amplicon (bp)** |
| --- | --- | --- | --- |
| aimR_fwd | acgaaaatcgccattcgccagggctgcagg  CCCTCATTGTGTTTAGGTAAAATAAG | 15 s at 98C  10 s at 65C  30 s at 72C  X35 cycles | 1371 |
| aimR_rev | Attcaattatttaa  TATTCTCACCTCCTTTCAAATTTG |  |  |
| aimX_fwd | Aaggaggtgagaata  TTAAATAATTGAATAGGTAATACATAATACTATC | 15 s at 98C  10 s at 57.7C  30 s at 72C  X35 cycles | 1238 |
| aimX_rev | Caactggtaatggtagcgaccggcgctcag  TAAATACGCTTCACAGTTTC |  |  |

Table S12. Phage mutants used in this study

| Phage mutant | Origin | Details |
| --- | --- | --- |
| Phi3TΔ*aimP* | Bruce et al, 2021 | Phi3T WT with the aimP knocked out, non-signal producing Phi3T |
| Phi3TΔ*aimRPX* | Bruce et al, 2021 | Phi3T WT with the aimRPX knocked out, non-signal producing and non-receiving Phi3T |
| WT Phi3T.*kan* | This paper | Wildtype Phi3T phage with a kanamycin cassette replacing *yokI* |
| Phi3T.SIIRGA.spc | This paper | Phi3T with the WT *aimR,* *aimP* genes and aimX promoter replaced with those of a SIIRGA phage, and spectinomycin cassette replacing *yokI* |
| Phi3T.SASRGA | This paper | Phi3T with the WT *aimR, aimP* genes and aimX promoter replaced with those of a SASRGA phage |
| Goe11*yokI*::*tet* | Jose Penades group | Goe11 with tetracycline cassette replacing *yokI* |

Table S13. sgRNAs, Phi3T WT homology template primers, and primers to amplify gblocks and the gblocks themselves used to engineer the Phi3T.SASRGA and Phi3T.SIIRGA arbitrium phage mutants and Phi3T.kan and Phi3T.SIIRGA.spc mutants.

| **Mutant/target** | **Primer or gblock sequence** | **Amplification program** | **Amplicon (bp)** |
| --- | --- | --- | --- |
| **SASRGA mutant**  sgRNA_aimR_F  sgRNA_aimR_R | **TACG**TCAAGCATTTCTTGGTGCAC  **AAAC**GTGCACCAAGAAATGCTTGA | n/a | n/a |
| HT_upstream_F | CGACTCACTATAGGGTCGACAAAATTGAGAAAGG  AGAAGCAAGAATACTTAATAACGGAACAATTGAG | 10 s at 98C  30 s at 64C  40 s at 72C  X35 cycles | 745 |
| HT_upstream_R | GAGACCGGTCTCCTGCTTTAATTTCAATTGTC  TCCCCCCTTTATTCAGTTATCACTTCCC |  |  |
| HT_d’stream_F | TAAAGCAGGAGACCGGTCTCGGGGGTTATTAGAA  TGAAAAGAGCATTAGGTAAAGCAATATCTTATGAAG | 10 s at 98C  30 s at 66C  40 s at 72C  X35 cycles | 750 |
| HT_d’stream_R | TATATTTTAGATGAAGATTATTTCTTAATCTAGAAAG  GCCTTGTCCCCTACATTTATCACAGGTTTACCAGTCAG |  |  |
| SASRGA_gblock_F | AAGTGATAACTGAATAAAGGGGGGAGACAATTGAAATTAAATTAAAGCAGATGAATCTTAAGCAGATGATTAAGAATG | 10 s at 98C  30 s at 68C  40 s at 72C  X35 cycles | 1514 |
| SASRGA_gblock_R | TAAGATATTGCTTTACCTAATGCTCTTTTCATTCTAATAA  AATAACCCCCTATTTGGTTTTATAAAATATAGGAACTTCC |  |  |
| SASRGA gblock | ATTAAAGCAGATGAATCTTAAGCAGATGATTAAGAATGAATGTGAAAAAGACAACCAGCTCGCAGCGAAACTCTCAAAAATAGCAGGGTACGAAAAGGTTAATGGTTTTTACAAATTCATCAACACCCCAGAGAAAGAAATGGACAACTTAGGCGGTTTAATTAATATTGTTAAGAGCTTGTTTCCGGATAATGAAGAGCAGCTTCTAAGTGACTACTTCTTATCATTGGATCCCAATAAAAAAAGCGCAAGACAGTCTGTCGAGTATGCAGATTTAAACCAATGGAATGCATTGACTGATAAGATCGTAAGCAATCTTTGCGAATCATCTAATTCAATAAGTCGTGAATGGGGACAGGTTTATTCCCTACATAGAAAACTGAATAATAATAAAATTTCTATAAATGAAGCGATCCGGGAAACTGGGAAATATAGAATTAAATCTCCTGAAATGTATTCATTTTCGAATATTATGATTATGTACGAATACTTGAAAATTGGAGAATTTGGCTTAATGAAAAGTACAGCTCAGTTTCTGGAGATTGACGAACTGTCTAATGGATTTATAAAAGATTCGTACAGTGGTCGAATTGAACTGTTAAAGGCCAATATAAGCTTAAATGATTATGAACTAGAAGAAACCCGAAAACATTGTAGCGCTGTAATTGAAGAATGCAATAATAACAGATTGATTGTATTTAGTTATTTAACACTTGGGAATACATACATTTTTGAAGATTATGCTAAAGCAAAACTATGCTATGAAAAAGGCTTGAACTTTGCAAAAGACAATAGCCATCATCATTATAAATTACGACTCGCACTTTGCTTTTTAGATAATGTCTGGGCGAGAGAAAACAAATGGGTAGATTTCGAGTCTCAAGAAATACCGGATATGATTGAAGCTGCTTTTTATTTGACTAATATCAAAGAAACTAAGAAAGCAGAAGATGTTATTAAAAAAATTGAAGAACATGATGTTCTGGATGATGATTTAGGGTTTCTTTATCACGTTAAGGGCTTGCTGTATAATGATATGTCCCATTTTCACGAGAGTATAAAGAAATTCAAAAAGTCAGGCGATAGGCTCTGTCTAAATCTACCTTTGATTGAATTGAAAAAGCATGGATACTCAGATGAAATATTAAATTTAATTGCGCTATAGTTTTCTTCACTTGAAAGGAGGTGAAAGAATGAAGAAATTTAATTGCGCGATTGTCATTTTACTAGCTTTAACTGTAGGATTTGTAAGTGGACAACAATCAGTCCAAACTGCTAACGGAGATATCACAGTGGCTTCAGCTAGCAGAGGAGCATAACACAACTAGGAATTTACATAGTCAACGGTTAGACGTTTGATCCAAAGGATCAGGCGTCTTTTCTAATTTAAAGGGGAAGTTCCTATATTTTATAAAACCAAATAGGGGGTTATT | n/a | 1433 |
| **SIIRGA mutant** |  |  |  |
| sgRNA for aimR F  sgRNA for aimR F | tacgAATTTGATGATTTACCCGAA  aaacTTCGGGTAAATCATCAAATT |  |  |
| HT_upstream_F | TACGACTCACTATAGGGTCGACGGCCAACGTTGCAGCTCCAGGTACGTTTG | 10 s at 98C  30 s at 68C  40 s at 72C  X35 cycles | 746 |
| HT_upstream_R | CTGCTTAAGATTCATCTGCTTTAATTTCAATTGTCTCCCC |  |  |
| HT_d’stream_F | AAAAAACAAGAACATGGGGGTTATTAGAATGAAAAG | 10 s at 98C  30 s at 65C  40 s at 72C  X35 cycles | 746 |
| HT_d’stream_R | AGATTATTTCTTAATCTAGAAAGGCCTTATATTTTCAAACCCTATCCATTC |  |  |
| SIIRGA_gblock_F | TTGAAATTAAAGCAGATGAATCTTAAGCAGATGATTAAG | 10 s at 98C  30 s at 63C  40 s at 72C  X35 cycles | 1455 |
| SIIRGA_gblock_R | ATTCTAATAACCCCCATGTTCTTGTTTTTTATATTATGAATTTC |  |  |
| SIIRGA gblock | ATGAATCTTAAGCAGATGATTAAGAATGAATGTGAAAAAGACAACCAGCTCGCAGCGAAACTCTCAAAAATAGCAGGGTACGAAAAGGTTAATGGTTTTTACAAATTCATCAACACCCCAGAGAAAGAAATGGACAACTTAGGCGGTTTAATTAATATTGTTAAAAGCTTATTTCCTGATAATGAAGAGCAACTTCTAAGCGATTATTTTTTATCATTGGATCCCAATAAAAAATGTGCTAGACAATCTGTTGAATATGCGGATTTAAATCAGTGGAATGCATTAACTGATAAAATAATTTTAAATTTATGCAATTCGAAAAATGCGACAAGTAAAGAATGGGGTAAGACCTATAATATACATAGAAAGTTAACAGAAAACAAGATATCCTTAACTGAAGCAATCAGGGAAACTGGAAAATGCAAAACAGCAGAAATGATATTTTTCTCAAATGCAATGTTAATGTATGAATACCTAAAGATCGGTGAATTTGGATTAATGAAAAGCACAGCAAAATTGCTGGATTTTCAAGGATTATCAGACGGTTACATAAAAGGTTTATACACCTCCAGAGTAAGCTTGTTGAAAGCTAATATAAGCTTCAATGAGAACAATTTAATTGAAGCAAGAAAATATTGTTTATATGCCACTGAAACTACGAACGTGGATAGGATTTGTTTTTTTGCATATTTAACAATCGGAAACTCTTTCATATTCGAAAATTTTGAAGAGGCCAAGCGATCATATATTAATGGTGCTAAGTATGCAAGCAACACAATTCATAAAGAGATGTTAGACGGAGCATTATGTTTTCTTGCAAGCTTTTGGAACAAAGAGAATTTATGGGTGAATTATGAATCACAGCACACTAAATACTTGCAATTGAGAGCATACCATCATATACGAAAAGGCGAAGTTGATAAAGCTAATGAGATTTTAAATGAGTTATCAATAAGAGAACAAGATGAGAATGAGATGGGATTTTATTTTTATTATAGAGGTTTAATATCTGTAGATAAATCTGATTTTTATAAATCTATACGCTGTTTCAAAAAATCAGATGACAAATATTCAGTTCAATTGCCCTTGATTGAACTTAAAAAAATGGGCGCGGACACAGAACTGTTAAGTCTTATTTCAATTTAGATTGATGCTATTGAAAGGGGGTGAGGTTGTTGAAAAAAACGATTTTAGGTGTAGCTATTATTGCGGCTCTGGCATTATCTTTTGTTGCAGGACAAAAATCAGTAAGCACTGCAGCTCCAAACGATGAAATTAGTGTAGCAAGCATTATTCGAGGGGCTTAAAAAATCACAACAGCGATACTACATAATCAATGGTTAGACGTTTGATCCAGTGGATCAGGCGTCTTTTCTAATTTTAAGAGAATGTTCCAGAAATTCATAATATAAAAAACAAGAACAT | n/a | 1425 |
| **Antibiotic mutants** |  |  |  |
| sgRNA for YokI F  sgRNA for YokI R | tacgGAAAGTAGGTTATGAGAATC  aaacGATTCTCATAACCTACTTTC | |  |
| upstreamF  upstreamR | TACGACTCACTATAGGGTCGACGGCCAACG  ATTGAAATCTTCTCTGATTTTG  GGGATTTCAACTGCAATCATTCTCCTTCCATTC | 10 s at 98C  30 s at 60C  40 s at 72C  X35 cycles | 1046 |
| **Kanamycin cassette**  F  R | TGGAAGGAGAATGATTGCAGTTGAAATCCCCTC  TCTTGAAAGCTCATATCAAAATGGTATGCGTTTTG | 10 s at 98C  30 s at 65C  40 s at 72C  X35 cycles | 915 |
| **Spectinomycin cassette**  F  R | TGGAAGGAGAATGATCGAATGGCGATTTTCGTTCG  TCTTGAAAGCTCATATCCCCCTATGCAAGGGTTTATTG | 10 s at 98C  30 s at 68C  40 s at 72C  X35 cycles | 1197 |
| D’streamF  d’streamR | CGCATACCATTTTGA  TATGAGCTTTCAAGATATCAATAATAAAATG  AGATTATTTCTTAATCTAGAAAGGCCTTAT  CAAGATGTATCTGAGTCAC | 10 s at 98C  30 s at 60C  40 s at 72C  X35 cycles | 1046 |

Table S14. Strain ID, species and origin used in the host range assays (BGSC = Bacillus Genetic Stock Centre)

| ID | host | origin |
| --- | --- | --- |
| 168 | *B.subtilis* | BGSC |
| 1723 | *B.subtilis IFO3022* | Eldar lab |
| 1726 | *B.subtilis IFO3215* | Eldar lab |
| 1729 | *B.subtilis IAM1232* | Eldar lab |
| 1732 | *B.subtilis spizizenii TU-B-10T* | Eldar lab |
| 1735 | *B.subtilis spizizenii DV1-B-1* | Eldar lab |
| 1738 | *B.subtilis AUSI98* | Eldar lab |
| 1741 | *B.subtilis RO-NN-1* | Eldar lab |
| 1771 | *B.atrophaeus 1942* | Eldar lab |
| 1774 | *B.subtilis "natto" IFO3335* | Eldar lab |
| 1777 | *B.subtilis "natto" IAM1163* | Eldar lab |
| 1780 | *B. subtilis "natto" NRRL B-14197 (=IFO1335)* | Eldar lab |
| 1802 | *B.mojavensis RS-A-2* | Eldar lab |
| 1808 | *B.vallismortis DV1-F-3 (=NRRL B-14890)* | Eldar lab |
| 3610 | *B.subtilis* | BGSC |
| 3738 | *B.subtilis* | Eldar lab |
| 6072 | *B.subtilis BEST7003* | Eldar lab |
| 7103 | *B. subtilis subtilis AUSI98 BGSC: 3A26* | Eldar lab |
| 7104 | *B. subtilis subtilis AUSI98 BGSC: 3A26* | Eldar lab |
| 7105 | *B.subtilis spizizenii ATCC 6633* | Eldar lab |
| 7108 | *B. subtilis inaquosorum KCTC 13429* | Eldar lab |
| 7111 | *B. subtilis spizizenii W23* | Eldar lab |
| 7112 | *B. subtilis Best7003 BTG1* | Eldar lab |
| 7113 | *B. subtilis ATCC 13952* | Eldar lab |
| 7114 | *B. subtilis DSM: 15029* | Eldar lab |
| Bt1 | *B. thuringiensis* | Penades/Marino lab |
| Bt2 | *B. thuringiensis* | Penades/Marino lab |
| Bt3 | *B. thuringiensis* | Penades/Marino lab |
| D6 | *B.subtilis168::D6* | Penades lab |
| Katmira | *B.subtilis* | Penades lab |
| ID | *host* | origin |
| 168 | *B.subtilis* | BGSC |
| 1723 | *B.subtilis IFO3022* | Eldar lab |
| 1726 | *B.subtilis IFO3215* | Eldar lab |
| 1729 | *B.subtilis IAM1232* | Eldar lab |
| 1732 | *B.subtilis spizizenii TU-B-10T* | Eldar lab |
| 1735 | *B.subtilis spizizenii DV1-B-1* | Eldar lab |
| 1738 | *B.subtilis AUSI98* | Eldar lab |
| 1741 | *B.subtilis RO-NN-1* | Eldar lab |
| 1771 | *B.atrophaeus 1942* | Eldar lab |
| 1774 | *B.subtilis "natto" IFO3335* | Eldar lab |
| 1777 | *B.subtilis "natto" IAM1163* | Eldar lab |
| 1780 | *B. subtilis "natto" NRRL B-14197 (=IFO1335)* | Eldar lab |
| 1802 | *B.mojavensis RS-A-2* | Eldar lab |
| 1808 | *B.vallismortis DV1-F-3 (=NRRL B-14890)* | Eldar lab |
| 3610 | *B.subtilis* | BGSC |
| 3738 | *B.subtilis* | Eldar lab |
| 6072 | *B.subtilis BEST7003* | Eldar lab |
| 7103 | *B. subtilis subtilis AUSI98 BGSC: 3A26* | Eldar lab |
| 7104 | *B. subtilis subtilis AUSI98 BGSC: 3A26* | Eldar lab |
| 7105 | *B.subtilis spizizenii ATCC 6633* | Eldar lab |
| 7108 | *B. subtilis inaquosorum KCTC 13429* | Eldar lab |
| 7111 | *B. subtilis spizizenii W23* | Eldar lab |
| 7112 | *B. subtilis Best7003 BTG1* | Eldar lab |
| 7113 | *B. subtilis ATCC 13952* | Eldar lab |
| 7114 | *B. subtilis DSM: 15029* | Eldar lab |
| Bt1 | *B. thuringiensis* | Penades/Marino lab |
| Bt2 | *B. thuringiensis* | Penades/Marino lab |
| Bt3 | *B. thuringiensis* | Penades/Marino lab |
| D6 | *B.subtilis168::D6* | Penades lab |
| Katmira | *B. subtilis* | Penades lab |

Table S15. Digital droplet PCR primers and conditions for quantifying the amount of phage produced during co-infections

| Primer name | Primer sequence | Target | PCR program | Amplicon (bp) |
| --- | --- | --- | --- | --- |
| Kanamycin-F | AATGGACAACCGGTGAGTGG | WT. Phi3T.kan | 5 m at 95C  30 s at 95C  60 s at 60C  (X40 cycles)  5 m 4C  5 m 90C  12 hrs 4C |  |
| Kanamycin-R | AAGCGGCCAATCTGATTCCA |  |  | 95 |
| Spectinomycin-F | TCGTCGTATCTGAACCATTGACA | Phi3T.SIIRGA |  |  |
| Spectinomycin-R | TTGGGAGGATGATTCCACGG |  |  | 195 |
| L6-7F  L6-7R | ACAAAGCAGCACATTCGCAA  CGGACATACCACCAACCTCC | Goe11.tet |  | 145 |
